# Magnetic Tweezers with Magnetic Flux Density Feedback Control

**DOI:** 10.1101/2020.12.03.410662

**Authors:** Waddah I. Moghram, Anton Kruger, Edward A. Sander, John C. Selby

## Abstract

In this work, we present a single-pole magnetic tweezers (MT) device designed for integration with substrate deformation tracking microscopy (DTM) and/or traction force microscopy (TFM) experiments intended to explore extracellular matrix rheology and human epidermal keratinocyte mechanobiology. Assembled from commercially available off-the-shelf electronics hardware and software, the MT device is amenable to replication in the basic biology laboratory. In contrast to conventional solenoid current-controlled MT devices, operation of this instrument is based on real-time feedback control of the magnetic flux density emanating from the blunt end of the needle core using a cascade control scheme and a digital proportional-integral-derivative (PID) controller. Algorithms that compensate for an apparent spatially non-uniform remnant magnetization of the needle core that develops during actuation are implemented into the feedback control scheme. Through optimization of PID gain scheduling, the MT device exhibits magnetization and demagnetization response times of less than 100 ms without overshoot over a wide range of magnetic flux density setpoints. Compared to current-based control, magnetic flux density-based control allows for more accurate and precise magnetic actuation forces by compensating for temperature increases within the needle core due to heat generated by the applied solenoid currents. Near field calibrations validate the ability of the MT device to actuate 4.5 μm-diameter superparamagnetic beads with forces up to 25 nN with maximum relative uncertainties of ±30% for beads positioned between 2.5 and 40 μm from the needle tip.

## I. INTRODUCTION

Despite advances in our understanding of the pathophysiology of congenital and acquired human blistering skin diseases, the biophysical mechanisms by which cell-cell and cell-matrix anchoring junctions endow epidermal keratinocytes with such an innate mechanical resilience are not completely understood^1^. Fortunately, during the past several decades, cell mechanics has been the subject of intense exploration amongst biologists, physicists, and engineers^2–4^. As such, numerous methodologies have been developed that can be used to apply mechanical loads and deformations to individual cells via cell-cell and cell-matrix anchoring junction proteins, including magnetic tweezers, optical tweezers, and magnetic twisting cytometry, to name a few^5–10^. Moreover, the well-established techniques of deformation tracking microscopy (DTM)^11,12^ and cell traction force microscopy (TFM)^13–16^ can be used to quantify substrate deformations and traction stresses present at cell-matrix junctions that individual cells and multicellular sheets use to attach to model biological surfaces^16,17^. As the overarching goal of this work, we set out to demonstrate how a scientific apparatus that *integrates* magnetic tweezers (MT) and substrate deformation tracking/traction force microscopy (DTM/TFM) can be employed to explore the mechanobiology of cell-cell and cell-matrix anchoring junctions within human epidermal keratinocytes cultured *in vitro* as a model for investigating skin fragility disorders. To date, the techniques of DTM and TFM have been widely incorporated into the biology laboratory due to the relative simplicity of the experimental setup and existence of open source computational machinery for analyzing image-based data^17–19^. Arguably, the implementation of MT devices is much more limited. Towards this end, in this first of two companion publications^20^, we present a detailed discourse on the concept, design, assembly, operation, and calibration of an MT device for ease of replication in the biology laboratory using commercially available off-the-shelf electronics, mechanical hardware, and computer software. In contrast to conventional MT devices that achieve magnetic actuation forces through feedback control of solenoid currents, our MT device is unique in that it is based on feedback control of the magnetic flux density emanating from the needle core. Potential advantages of this control configuration are demonstrated herein.

## II. MAGNETIC TWEEZERS – CONCEPT

In theory, operation of a single-pole MT device is based on the principle that an electrical current, *I*, applied to a solenoid coil surrounding a soft ferromagnetic core will induce a magnetic field in the core that emanates from both the blunt end of the core and the needle tip (**Fig. 1(a)**)^21^. The magnetic field within the core, ***B***_*c*_, at any point in time, *t*, is a function of *I*; the relative permeability of the core, *μ*_*c*_; the saturation magnetization of the core, 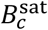; and the coercivity of the core, *H_c_*. Note that *H_c_* is closely related to the remnant magnetization of the core, 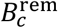, the magnitude of which depends on the material’s magnetization history. Here, *μ*_*c*_(*T*), 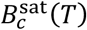, and 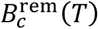 can all be regarded as intrinsic material properties of the core that all have a temperature dependence, denoted by the variable, *T*. If a superparamagnetic bead is exposed to the magnetic field present in front of the needle tip, ***B***_tip_, the MT will impose a magnetic force on the bead, **F**_*MT*_, that is a function of bead diameter, *d*; the relative permeability of the bead, *μ*_*b*_; and gradients present in the magnetic field, ▽***B***_tip_; given by the relationship:

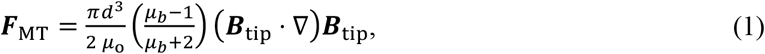

where *μ*_0_ is the permeability of free space^22^. Here Equation (1) assumes that the magnetization of the superparamagnetic bead does not saturate in response to ***B***_tip_. Magnitudes of the field gradients near the tip are dependent on the physical geometry of the needle, magnetic properties of the core, and the strength of the magnetic field generated by the solenoid coil. From an experimental standpoint, the magnetic actuation force generated by an MT on a superparamagnetic bead can be considered to be a function of the spatial positioning vector of the centroid of the bead with respect to the needle tip, **δ** and of ***B***_tip_, where the latter is a function of *I*, *μ*_*c*_(*T*), 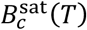, and 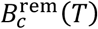, or

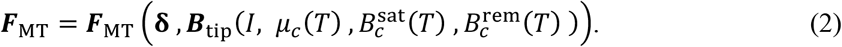

**FIG. 1.**
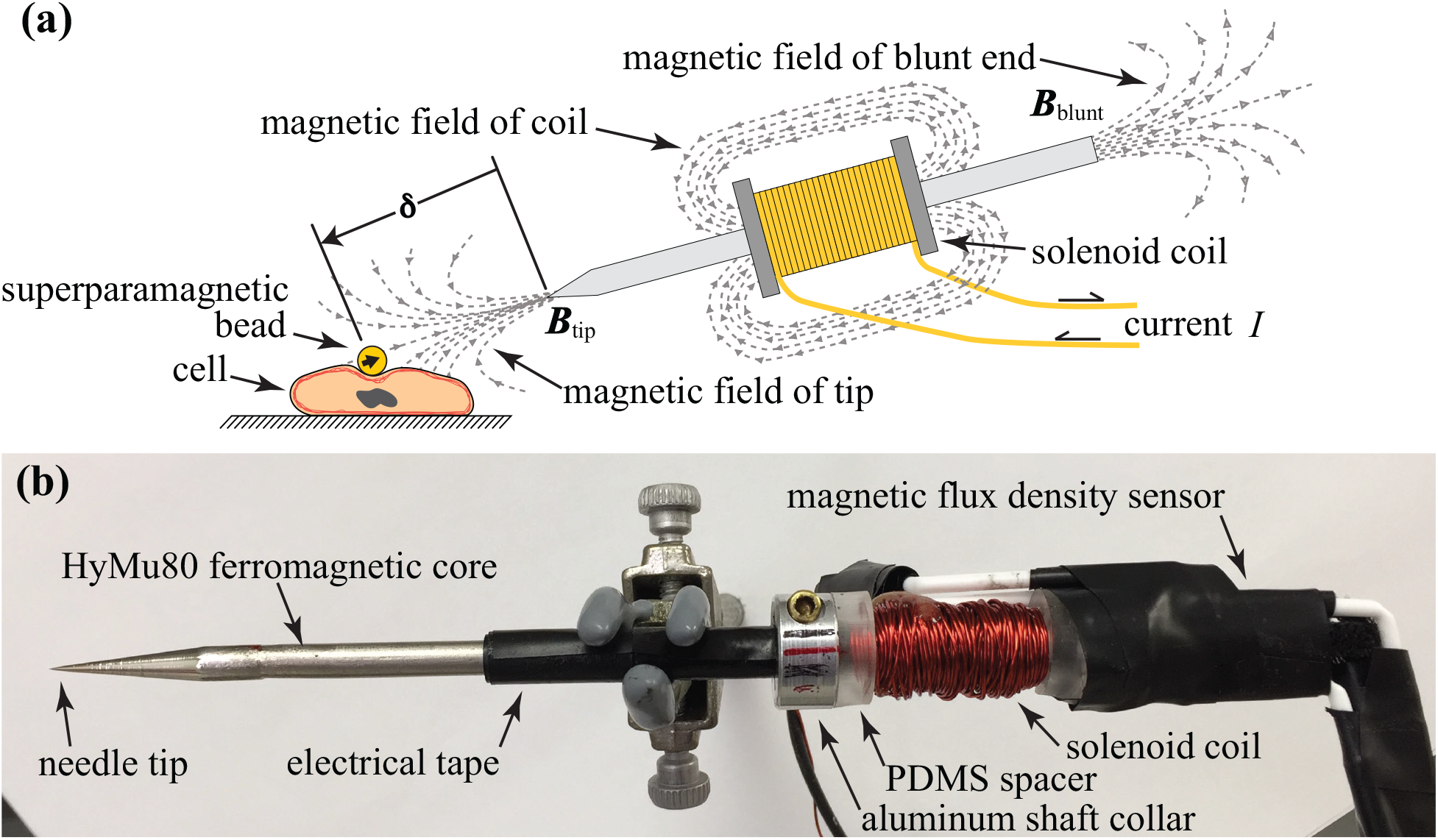
**(a)** A conceptual schematic showing the principle of magnetic force actuation in an MT device. The magnetic actuation force, **F**_*MT*_, applied to a superparamagnetic bead attached to a cell or biologic substrate (denoted by the black block arrow in the bead) is a function of gradients in the magnetic field emanating from the needle tip, ***B***_tip_, and the spatial positioning of the bead relative to the needle tip, **δ**. **(b)** Photograph of the actual MT needle assembly used in this work.

Ideally, the most accurate method for controlling **F**_MT_ in an MT experiment would be through feedback control of ***B***_tip_. However, placement of one or more magnetic flux density sensors within the proximity of the needle tip to provide feedback measurements of ***B***_tip_ is not practical for most applications. As such, conventional MT designs are based on real-time feedback control of *I* to *indirectly* control ***B***_tip_, and thus ***F***_MT_ for a given **δ**^22,23^. Using image-based particle tracking, it has also been shown possible to monitor **δ** in real-time such that closed-loop feedback control of ***F***_MT_ can be achieved by adjusting either *I* or **δ**, the latter accomplished through rapid spatial positioning of the needle tip using a motorized micromanipulator^23^. Despite these remarkable advances in MT devices, it is important to recognize that all core materials develop remnant magnetization after exposure to an applied magnetic field. Additionally, temperature increases in the needle core due to heat generated from current flow in the solenoid coil will lead to changes in *μ*_*c*_(*T*), 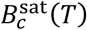, and 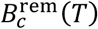. Collectively, thermally-induced fluctuations within the electron structure of the core material can ultimately *oppose* magnetization by an applied field^24^. All of these effects can lead to erroneous control of ***F***_MT_ if one utilizes only solenoid currents, *I*, to control ***B***_tip_ while disregarding the thermal and magnetic history of the core. To nullify these effects, many research groups have employed various combinations of in-operation degaussing current bursts^23,25,26^, compensation currents based on magnetization history^23^, and/or adjunctive cooling systems^27–29^. As an alternative and much less widely adopted approach^30–32^, we demonstrate control of ***F***_MT_ based on feedback control of the magnetic field present at the blunt end of the needle core, ***B***_blunt_, or conceptually,

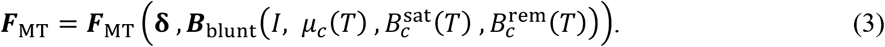

For ease of review, a comprehensive list of abbreviations and symbols can be found in the **Supplementary Material**.

## III. DESIGN

### A. Hardware

Much of the design of our single-pole MT device is modeled on the work of Kollmannsberger and Fabry^23^, modified to implement a magnetic flux density-based control system. As detailed in **Fig. 2**, the instrumentation in our integrated MT-DTM/TFM setup includes a Nikon Eclipse Ti-E inverted microscope (Nikon, Melville, NY) operated by an HP Z72 Workstation (referred to as *host* computer) that runs Nikon Elements AR 4.51 software on a 64-bit Windows^®^ operating system. The inverted microscope is equipped with both standard differential interference contrast (DIC) and epifluorescence imaging modalities, the latter of which employs a Texas Red bandpass filter cube set (560/40 nm excitation filter, 595 nm dichroic mirror, 630/60 nm emission filter). Integrated MT-DTM/TFM experiments were conducted using Nikon Plan Apo Lambda 10X (0.30 NA) or CFI Plan Apo VC 20X (0.75 NA) air objective lenses that were optionally coupled with a 1.5X magnifier. MT calibrations (see Sec. V) used a Nikon CFI S Plan Fluor ELWD 40X (0.60 NA) objective lens. A PCO.EDGE 4.2 sCMOS camera (PCO-Tech, Wilmington, DE) was used to capture both DIC and epifluorescence images at a standard frame rate of 40 frames/second (fps) and a resolution of 2048×2044 pixels, with the option of increasing frame rates to 100 fps for smaller defined regions of interest (ROIs).

**FIG. 2.**
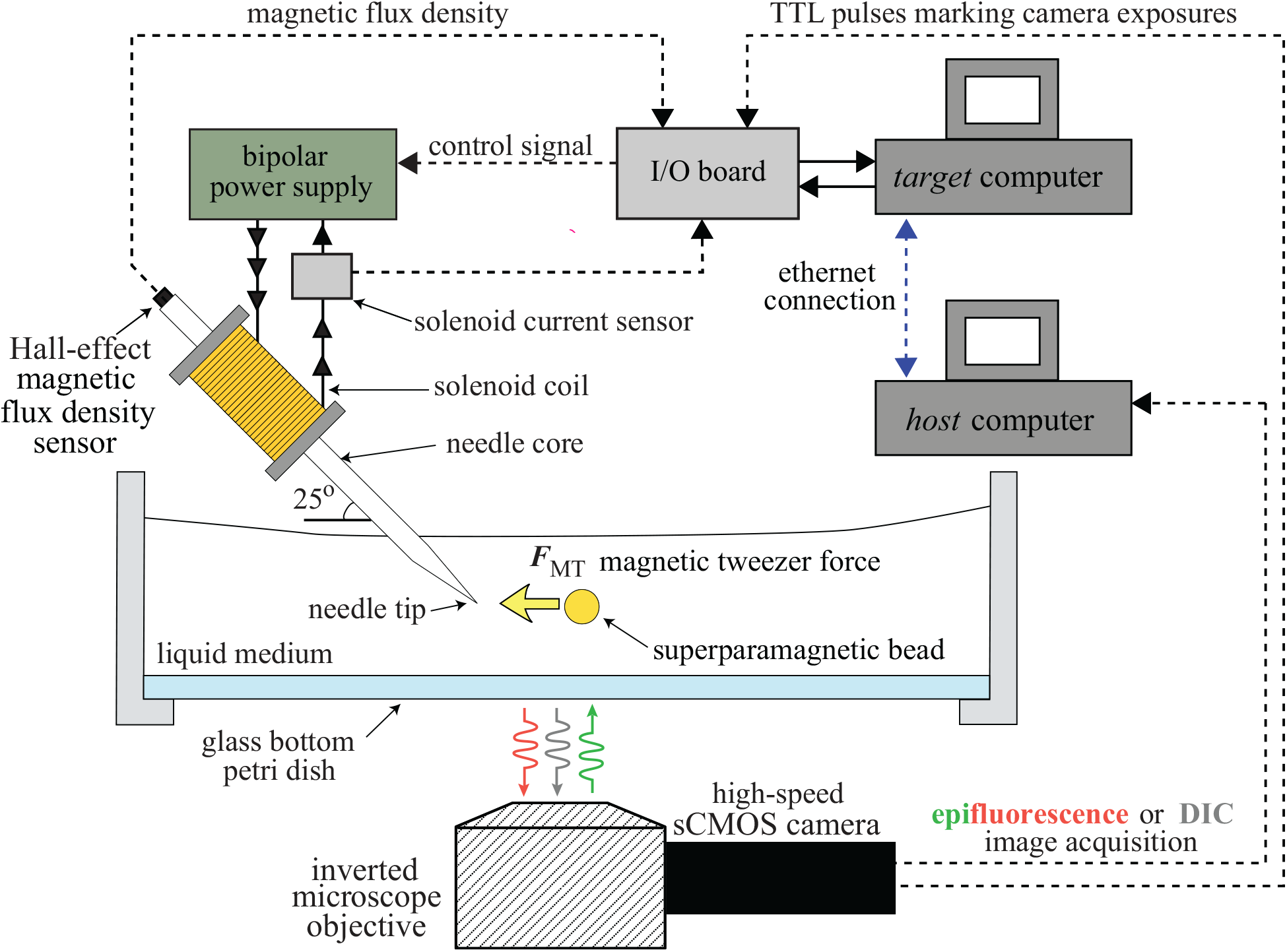
Schematic exhibits the key instrumentation used in an MT device configured for real-time feedback control of *B*_blunt_ (see text for details). Magnetic actuation force sequences and fast time-lapse acquisition of epifluorescence or DIC images are synchronized via an ethernet connection between the *host* and *target* computers. TTL pulses marking exposures of the sCMOS camera allow temporal alignment of data derived from captured images and magnetic flux density measurements. During operation, the needle core is positioned at an angle of 25° with the horizontal, designated as the *standard configuration*.

In addition to the Nikon-supplied 12-bit NI PCI-6713 PCI data acquisition (DAQ) board used to control the microscope, the *host* computer has also been outfitted with a 16-bit NI PCIe-6321 DAQ board and an SCB-68A connector block operated by an NI LabVIEW™ 2015 software package that includes a Real-Time Module (National Instruments, Austin, TX). The *host* computer controls the microscope, acquires fast time-lapse images, and deploys custom-made LabVIEW™ programs via an ethernet connection to a second *target* computer that controls the MT device. The *target* computer is a Dell Optiplex 9010 equipped with a 16-bit NI PCIe-6341 DAQ board and an SCB-68A connector block that is used to acquire analog input measurements (magnetic flux density, solenoid current, and transistor-transistor logic (TTL) pulses marking exposures of the sCMOS camera) while generating analog output voltages to control actuation of the MT device in real-time. The *target* computer is operated by LabVIEW™ Real-Time 15.0.1 deterministic operating system (National Instruments, Austin, TX).

The MT is constructed from a cylindrical ASTM A753-08 Alloy Type 4 soft ferromagnetic core (Scientific Alloys, Westerly, RI), 4.76 mm in diameter and ~165 mm long, machined with one blunt end orthogonal to the longitudinal axis of the core and the other end tapered at an angle of 7.5° from the longitudinal axis (**Fig. 1(b)**). The core was hydrogen annealed at 1175°C for 4 to 6 hours, followed by controlled cooling to 370°C at a rate of 175-315°C/hour (Vac-Met Inc., Warren, MI). After annealing, the tapered tip was hand-sharpened to a needle point with a radius of curvature of ~10 *μ*m using a series of 1200-, 4000-, 6000-, and 8000-grit polishing cloths (3M, St. Paul, MN). Roughly 1 cm away from the blunt end, the core is wrapped with a single layer of insulating PVC electrical tape (3M) flanked by two 2024 aluminum shaft collars spaced ~35 mm apart and fixed in place with brass socket head cap screws. A solenoid coil consisting ~250 turns of 24 AWG enameled copper wire (Remington Industries, Johnsburg, IL) is wound between the collars and insulated from the aluminum using Sylgard^®^ 184 polydimethylsiloxane (PDMS) spacers (Dow Corning, Midland, MI) fabricated in-house. A Honeywell SS-495A1 Hall-effect magnetic flux density sensor (Honeywell, Charlotte, NC) and its wiring harness are affixed to the blunt end of the needle with the sensing loop positioned concentrically with the cylindrical needle core using a combination of 2- to 3-mm-wide Velcro strips (3M) epoxied to the sides of the cylindrical core, ~2.5-mm diameter polypropylene guide rods, and PVC electrical tape. The MT is mounted on an MHW-3 three-axis water hydraulic manual micromanipulator (Narishige, Amityville, NY) that was fitted with an aluminum adaptor arm fabricated in-house to extend the reach of the needle tip over the microscope stage. The magnetic flux density sensor is powered by a 5VDC linear power supply (Bel Power Solutions, Santa Clara, CA). The solenoid coil is energized by a Kepco Model BOP 100-4DL bipolar power supply (Kepco, Flushing NY) controlled by analog voltages output from the DAQ input/output (I/O) board in the *target* computer. Solenoid current measurements are acquired by monitoring the voltage drop across a 1 Ω ±1%/50W power resistor (Vishay, Malvern, PA) configured in series with the solenoid coil and confined in a separate electrical housing. Magnetic flux density sensor output voltages, sCMOS camera exposure TTL pulses, and solenoid current measurements are all acquired by the DAQ Input/Output (I/O) board of the *target* computer. All inputs measurements are sampled at 30 kHz. Photographs of the apparatus can be found in **Figs. S1** and **S2** of the **Supplementary Material**.

### B. Magnetic flux density-control scheme

As a surrogate for ***B***_tip_, operation of our MT device is based on control of the magnetic flux density emanating in the axial direction from the blunt end of the needle core, a quantity we define as *B*_blunt_. Our feedback control routine utilizes a cascade configuration with a *master* or *outer* loop that uses a 1.0 kHz digital proportional-integral-derivative (PID) controller within the *target* computer for setpoint control of *B*_blunt_, and a faster *slave* or *inner* analog loop operating within the bipolar power supply for setpoint control of *I*. In the *slave* loop, the bipolar power supply drives *I* to the setpoint corresponding to the analog voltage signal output from the *master* loop. The bipolar power supply is specifically designed for powering inductive loads with typical response times of ~220 to 280 μs at 1.7 kHz for a 2-mH nominal load. These specifications are similar to those reported for other MT devices^23,24,33^. The master loop uses consecutive 30-sample averages of *B*_blunt_ measurements to update analog output control voltages that are subsequently passed to the bipolar power supply (i.e., *slave* loop) (see **Fig. 2)**.

## IV. OPERATION

### A. Degaussing and magnetic properties of the needle core

Prior to any initial actuation of the MT device, a degaussing current is applied to the solenoid to demagnetize the needle as part of our standard operating protocol. Specifically, the degaussing routine is composed of 3 consecutive applications of a 60 Hz current waveform, *I*(*t*)_dmag_, defined as *I*(*t*)_dmag_ = *I*_0_ sin(120 *π t*) exp(−*t*^2^), where the initial current, *I*_0_ = 4 A, and *t* spans from 0 to 5 s. To facilitate tuning of the PID control parameters for the *master* loop, we first characterized the magnetic behavior of the HyMu80 core. The longitudinal axis of the needle core was inclined at an angle of 25° from the horizontal (defined as the *standard* MT device configuration), and a second magnetic flux density sensor (Honeywell SS-495A1) was vertically oriented with the center of its sensing loop positioned ~100 μm lateral to the needle tip within the same horizontal plane. This second sensor was used to measure the magnetic flux density at this location/position, an experimental quantity defined here as *B*_tip_. Following degaussing of the core, the current in the solenoid was incrementally looped with the device configured for conventional feedback control of *I* from 0 A to −4 A, then from −4 A to 0 A, then from 0 A to 4 A, and finally from 4 A back to 0 A over a period of 30 s while recording *B*_blunt_ and *B*_tip_. The results of this characterization are shown in **Fig. 3**. The saturation flux density for *B*_blunt_ was approximately ±200 Gs with a linear response between ±150 Gs, and minimal hysteresis due to hydrogen annealing of the needle core. *B*_tip_ was observed to exhibit a much more exaggerated hysteresis loop in response to the applied solenoid current loop, with a remnant magnetic flux density of roughly −1.95 Gs at the end of the 0 A to −4 A to 0 A sequence compared to remnant magnetic flux density of 0.75 Gs measured for *B*_blunt_. Note that the signs of *B*_tip_ and *B*_blunt_ are opposite, consistent with the physical orientation of magnetic field lines that are generated during actuation of the MT device.

**FIG. 3.**
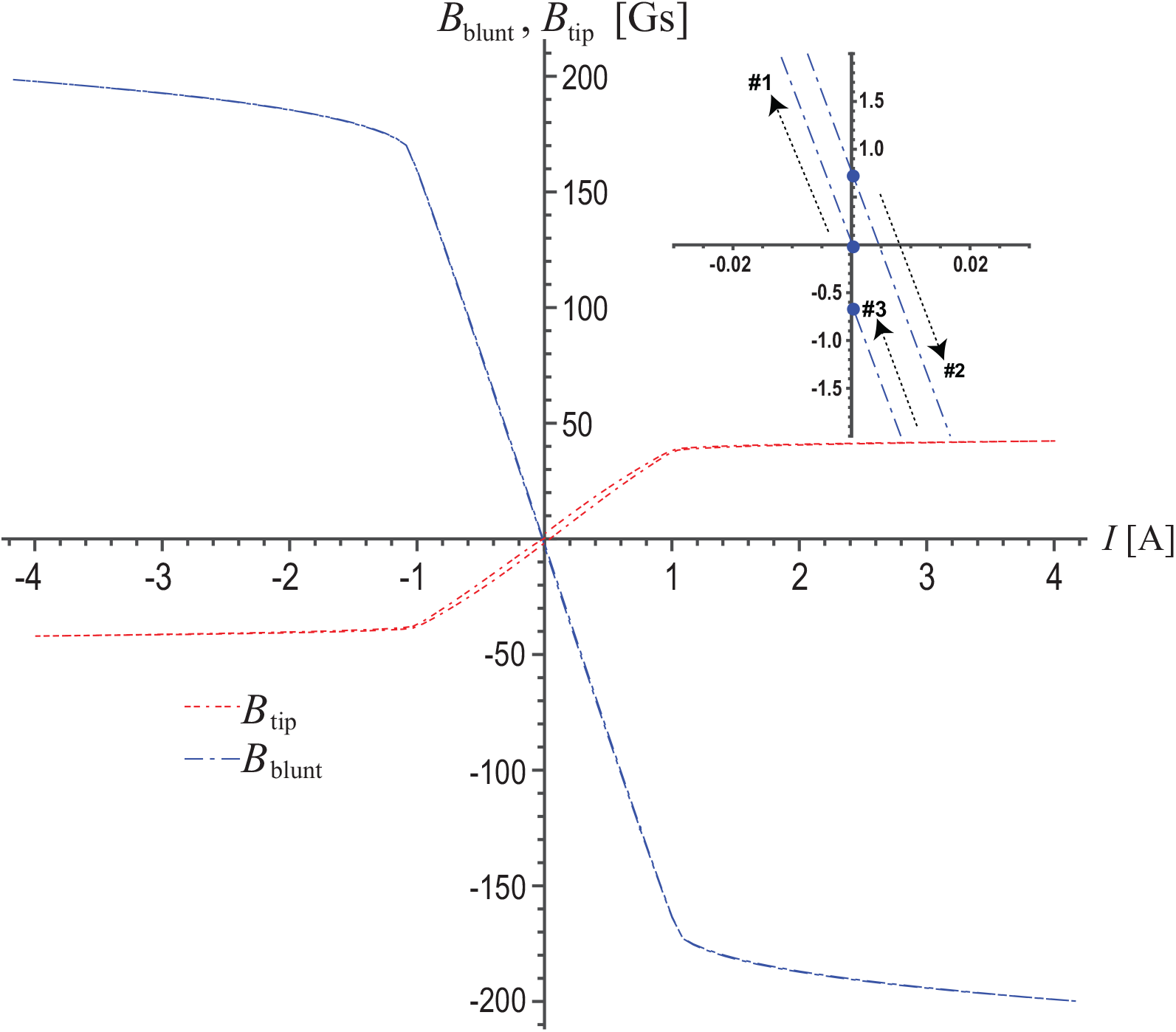
Characterization of the magnetic response of the HyMu80 alloy MT needle core. Directions of the solenoid current loop sequence are denoted by black dashed arrows in the figure inset. A blue dashed line represents *B*_blunt_ measurements and a red dashed line represents *B*_tip_ measurements. Inset shows an expanded view of the origin where *B*_tip_ measurements have been omitted to demonstrate how minimal magnetic hysteresis is observed in *B*_blunt_.

### B. Actuation response times

To demonstrate the operating characteristics of the MT device under *B*_blunt_-control, a typical OFF/ON waveform sequence of 0.25 Hz is used, as shown in **Fig. 4(a)**. The output of the magnetic flux density sensor following completion of the initial degaussing routine is set as the zero magnetic flux density reference point, or 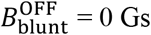. PID gain schedules in LabVIEW™ were empirically derived to achieve the widest range of controllable 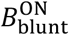 settings (i.e., ±185 Gs) to maximize the range of magnetic actuation forces, ***F***_MT_. PID gain settings were also set to minimize response times required to achieve these *B*_blunt_ OFF and ON setpoints (denoted as 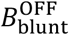 and 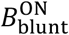, respectively) while explicitly avoiding overshoot (see **Figs. 4(b)** and **4(c)**). Overshoot in 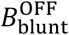 and 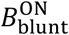, if present in an actual MT experiment, would lead to overshoot of ***F***_MT_. The D-term for all PID gains was found to be negligible for all operating conditions. With PID gain scheduling, our MT device was able to reliably control 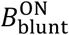 and 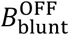 setpoints to within ±0.005 Gs (for time-averaged data) for magnetic flux density setpoints ranging from 0 Gs to ±185 Gs. Collectively, response times over the entire range of 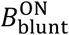 setpoints were ≤75 ms (magnetization or loading response times), while response times to achieve 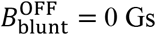 were ≤100 ms (demagnetization or unloading response times). In comparison to the ~5 ms response times of a recently published *B*_blunt_-controlled MT device^32^ and the 1 to 5 ms-response times of most *I*-control MT systems^23,34^, the response time of our *B*_blunt_-controlled device is predictably slower, a consequence of the cascade feedback control scheme and the 1 kHz digital PID control loop. However, we note that with upgraded data acquisition AI/AO hardware, it would be possible to sample *B*_blunt_ at higher rates (~90 kS/s) while increasing the frequency of our digital PID control loop (~3 kHz), likely enabling faster magnetization/demagnetization response times. More importantly, with programmable digital PID gain scheduling, our cascade control loop enables optimization of response times without overshoot in flux density setpoints *across the entire magnetic induction response of the core material*, including within the domain of non-linear magnetic saturation. In contrast, given a specified set of analog circuit components, hardware-based feedback systems for *B*_blunt_-control would be limited to optimized flux density responses within smaller defined domains of the overall magnetic induction response. Consequently, hardware-based feedback systems for *B*_blunt_-control are more likely to exhibit overshoot in magnetic flux density setpoints during operation^32^.

**FIG. 4.**
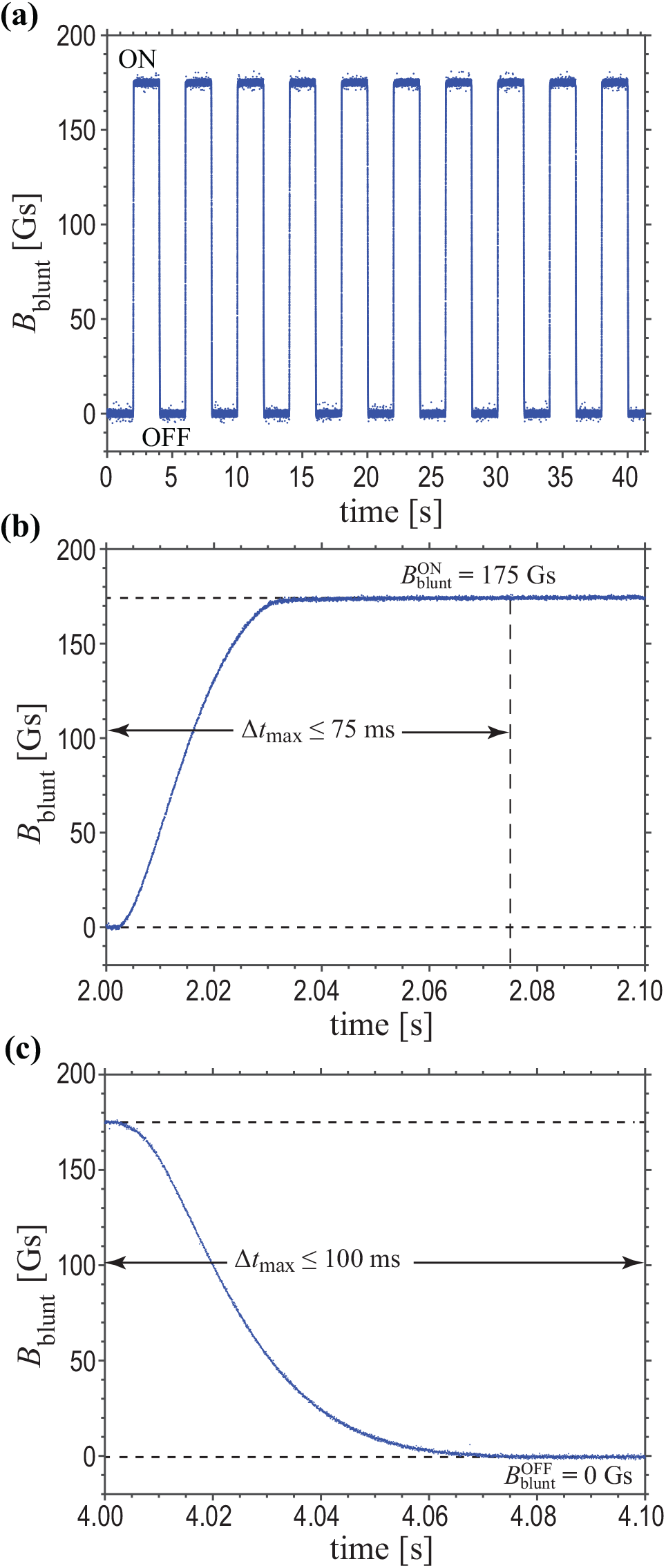
Operating characteristics of the MT device. A typical 10-cycle *B*_blunt_-control OFF/ON waveform sequence of 0/175 Gs at 0.25 Hz is depicted in **(a)**. Magnified view of the first ON segment of the magnetic flux density waveform is shown in **(b)**, where the magnetic actuation setpoint, 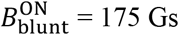 is reached within ~40 ms. Similarly, **(c)** depicts a magnified view of the first OFF segment, where the demagnetization setpoint 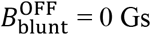 is reached within ~80 ms. Both ON and OFF *B*_blunt_ setpoints are achieved without overshoot.

### C. Comparison to conventional current-based feedback control

In *B*_blunt_-control mode, our MT instrument inherently achieves the prescribed *B*_blunt_ ON and OFF setpoints in a manner that (i) overcomes thermal effects that alter the magnetization properties of the core and (ii) negates the remnant magnetization that develops within the core following extinction of the field that was generated during the ON portion of the actuation sequence. **Fig. 5** showcases these two major benefits of magnetic flux density-based feedback control as opposed to current-based feedback control. Note how over the sequential ON segments of a typical 10-cycle 0/175 Gs OFF/ON 0.125 Hz waveform sequence, 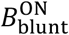 is maintained precisely at 175.0 Gs (**Fig. 5(a)**). However, due to heating of the core from the solenoid currents, the coil currents, *I*, required to achieve this magnetization ON setpoint increase in magnitude with each actuation cycle. Conversely, if a similar 10-cycle OFF/ON waveform is conducted using *I*-control with an ON setpoint of −3.0 A (higher current intentionally selected to illustrate thermal effects), the corresponding setpoints of 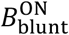 diminish as cycle number increases (**Fig. 5(b)**). Admittedly, in first 80 seconds in the *I*-control MT experiment, the decrease in 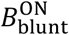 is small and would be associated with negligible variations in ***F***_MT_. However, for experiments of a much longer duration or larger currents where heating of the core becomes more pronounced, magnetic flux density-based feedback control would provide more reliable cycle-to-cycle control of ***F***_MT_. In **Fig. 5(c)**, notice how the solenoid current for *B*_blunt_-control is positive during the 0 Gs OFF setpoints the for cycles #2 through #10 **(Fig. 5(c)**). In other words, in the *B*_blunt_-control scheme, a small reverse current is automatically generated to null the small remnant magnetization that develops within the core following the magnetization of the ON segment. In contrast, in *I*-control mode, *B*_blunt_ OFF setpoints do *not* return to 0 Gs for cycles #2 through #10 as *I* returns to 0 A **(Fig. 5(d)**).

**FIG. 5.**
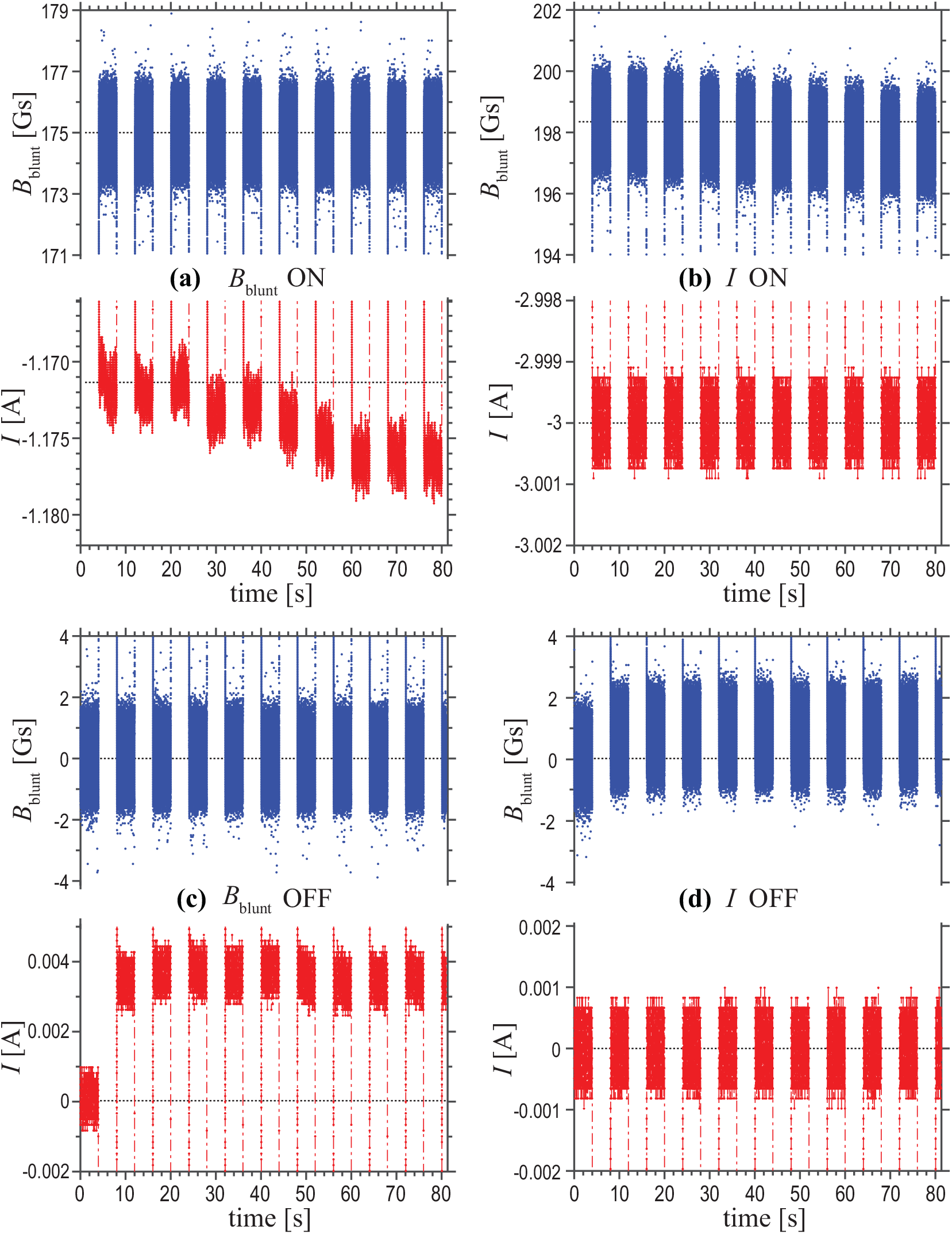
Comparison of magnetic flux density-control versus solenoid current-control of the MT device. A 10-cycle *B*_blunt_-control OFF/ON waveform sequence of 0/175 Gs at 0.125 Hz is depicted in **(a)** and **(c)**. In **(a)**, solenoid currents of increasing magnitude are required to maintain 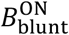 at 175 Gs with increasing cycle number, while in **(c)**, a small positive current is required to maintain 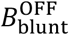 at 0 Gs for cycles #2 through #10. A 10-cycle *I*-control OFF/ON waveform sequence of 0/−3.0 A at 0.125 Hz is depicted in **(b)** and **(d)**. As observed in **(b)**, values of 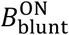 drift downward with successive cycles due to heating of the core. In **(d)**, 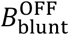 values do not return to 0 Gs after the first actuation cycle despite zero solenoid currents.

### D. Non-uniform remnant magnetization of the needle

To investigate whether *B*_blunt_-control can serve as a reproducible surrogate for controlling ***B***_tip_, we performed a series of experiments in which the MT device was oriented in the standard configuration and then actuated using OFF/ON *B*_blunt_-controlled 0.25-Hz waveform sequences while simultaneously measuring the magnetic flux density, *B*_tip_, at a spatial location ~100 *μ*m from the needle tip using a second vertically oriented Hall effect sensor (see **Fig. 6(a)** and **Sec. IV.A**). 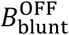 was set to 0 Gs for all test waveforms, and 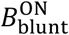 setpoints spanned from 0 Gs to 185 Gs. A typical sequence for a 0/175 Gs *B*_blunt_ OFF/ON waveform is shown in **Fig. 6(b)**. As previously observed, 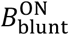 and 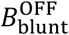 setpoints exhibit minimal to no cycle-to-cycle variation. However, for the second and for all subsequent OFF setpoints of the *B*_blunt_-control sequence, the value of 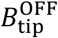 *deviates* from its initial (cycle #1) value by ~2 Gs for cycles #2 through #10, whereas 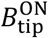 values showed no significant cycle-to-cycle variation. To explore this more quantitatively, we measured the variation in ON and OFF setpoint offsets of *B*_blunt_ and *B*_tip_ as a function of 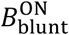. Here, 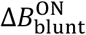 is defined as the difference in the time-averaged ON setpoints observed between cycle #10 and cycle #1, as measured by the magnetic flux density sensor positioned at the blunt end of the core. Similarly, 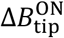 is defined as the difference in the time-averaged ON setpoints observed between cycle #10 and cycle #1, as measured by the magnetic flux density sensor positioned ~100 *μ*m from the needle tip. 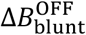 and 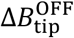 are defined in a consistent manner. To exclude transient effects, data from the first 500 ms and the last 500 ms of each 2 s ON or OFF setpoint duration were not included in the calculated time averages.

**FIG. 6.**
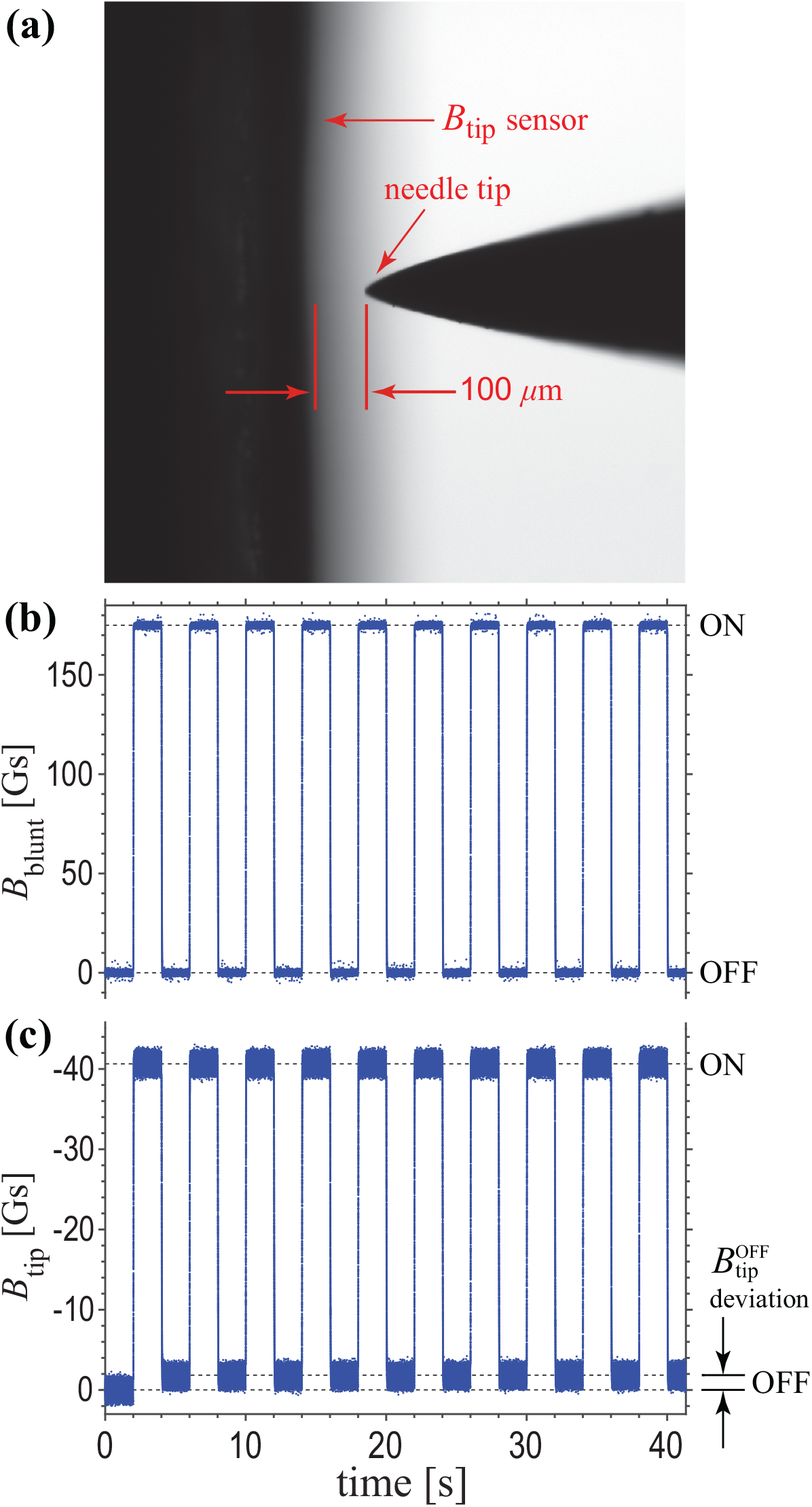
With the MT device maintained in the standard configuration, a second magnetic flux density sensor was vertically oriented and positioned ~100 *μ*m in front of the needle tip to quantify *B*_tip_, as shown in **(a)**. The MT device was then actuated with a 10-cycle 0.25 Hz *B*_blunt_-control 0/175 Gs OFF/ON waveform sequence **(b)** while simultaneously recording measurements at *B*_tip_. As can be seen in **(c)**, 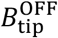 values do not return to 0 Gs for following the first actuation cycle.

**Fig. 7** shows the aggregate data from numerous replicates of these OFF/ON 0.25-Hz waveform experiments across a range of 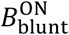 setpoints. Adjunctive measurements of *B*_tip_ that were collected by positioning the magnetic flux density sensor 200 *μ*m and 500 *μ*m from the needle tip demonstrated qualitatively similar results (data not shown). Clearly, the data justify the use of *B*_blunt_-control as a surrogate for control of *B*_tip_ during magnetic actuation of the needle core. As shown in **Fig. 7(a)**, magnetic flux density-based control is associated with 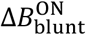 variations of ±5 mGs while 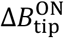 variations are repeatably ±0.2 Gs over the 10 cycles of operation (see **Fig. 7(b)**). However, despite precise and accurate control of 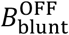 setpoints as hallmarked by 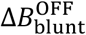 variations of ±5 mGs (see **Fig. 7(c)**), the magnitude of 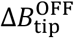 monotonically increases as a function of the 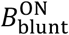 setpoint (see **Fig. 7(d)**). In other words, even though we had nullified the remnant magnetization present at the blunt end of the core following actuation of the MT device, the needle tip still maintained a finite remnant magnetization at OFF setpoints. Interestingly, evidence of non-uniform remnant magnetization of the needle core in a *B*_blunt_-controlled MT device was also observed in the stepwise force characterization presented by Kah et al.^32^, where 5.09 *μ*m-diameter superparamagnetic beads were subject to a finite MT force of ~0.23 nN at a bead-tip distance of ~30 *μ*m despite controlling for a 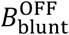 setpoint of 0 Gs.

**FIG. 7.**
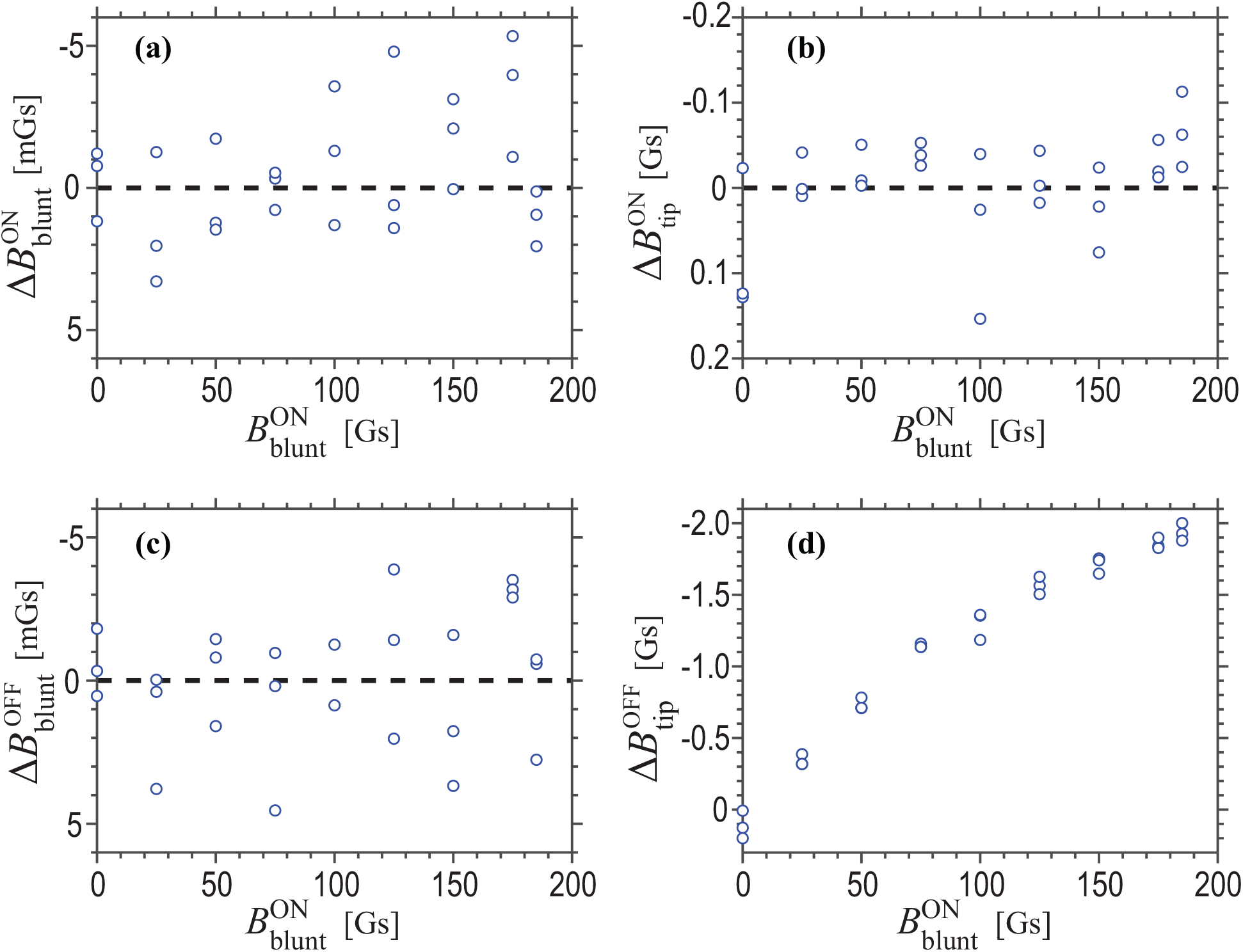
With the MT device in the standard configuration and a second magnetic flux density sensor used to quantify *B*_tip_, the MT device was actuated with 10-cycle 0.25 Hz *B*_blunt_-control waveforms with 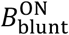 setpoints ranging from 0 Gs to 185 Gs, and 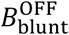 setpoints set to 0 Gs. Differences in measured *B*_blunt_ and *B*_tip_ ON/OFF setpoints between cycle #10 and cycle #1, defined as 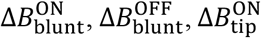, and 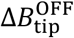, respectively, are plotted as functions of 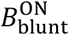 setpoints in **(a)**, **(b)**, **(c)**, and **(d)**. The magnitude of 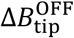 monotonically increases with increasing 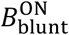, suggestive of an apparent spatially non-uniform remnant magnetization of the needle core.

Spatially non-uniform remnant magnetization was also observed in other MT needle prototypes developed during the course of this work. Specifically, less pronounced differences in remnant magnetization between the needle tip and the blunt end of the core were observed for needles with a 15° taper compared to the 7.5° taper of the needle shown in **Fig. 6(a)**. As such, we speculate that these observations might be attributed to differences in the length scale of magnetic domains within the HyMu80 Ni/Fe alloy relative to the length scales that define the bulk geometry of the core and its transition to the needle tip. Previous numerical modeling of MT devices has suggested that the magnitude of the magnetic field that develops within a soft ferromagnetic core in response to an energized solenoid varies along the longitudinal axis of the core as a function of core radius^33^. Assuming this to be true, it seems only logical to conclude that a spatially non-uniform remnant magnetization would be present upon removal of the applied magnetic field. Given the applied nature of this work, however, the true physical underpinnings of the apparent non-uniform remnant magnetization of the needle core were not further investigated.

### E. Compensation for null magnetic force actuation

Even after degaussing the needle core, the MT device was observed to exert forces on superparamagnetic beads suspended in glycerol and positioned in close proximity to the tip, i.e., ***B***_tip_ ≠ 0 Gs. A phenomenon that others have also observed^35^, we postulate that the small magnetic field present at the needle tip following degaussing is due to either a distortion of the earth’s magnetic field or a permanent magnetization of the needle tip. Collectively then, based on core characterization and *B*_blunt_-control experiments (see **Secs. IV.A** and **IV.D**), we identified two sources of magnetization that could lead to non-zero ***B***_tip_ fields, and thus non-zero ***F***_MT_, despite controlling *B*_blunt_ to output 0 Gs. The first source is referred to as *permanent* tip magnetization, i.e., the presence of a non-zero ***B***_tip_ despite degaussing of the needle core. The second source is referred to as the *remnant* tip magnetization, a term used to describe the apparent non-zero ***B***_tip_ that exists despite controlling *B*_blunt_ to output 0 Gs following any magnetization of the core.

In order to maintain 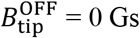 and hence *null* magnetic actuation forces (***F***_MT_ = 0 nN), additional corrective schemes were required to automate control of 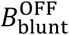 for actuation waveforms with 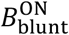 setpoints in the range of 0 to 185 Gs. To develop these corrective schemes, 4.5 *μ*m-diameter Dynabeads M-450 Tosylactivated superparamagnetic beads (Invitrogen, Waltham, MA) were suspended in pure glycerol (Sigma Aldrich, St. Louis, MO) and dispensed onto 50 mm-diameter petri dishes with 35 mm-diameter No. 1.5 coverslip glass bottoms (MatTek, Ashland, MA) subject to room temperature and ambient humidity of the laboratory (~10% to 50% relative humidity). The needle core was placed in the standard configuration and a degaussing routine was applied to the MT device. The needle tip was then positioned at δ = ~5 *μ*m from the bead, where δ denotes the magnitude of the positional vector, **δ**, as previously described (see **Fig. 1(a)**). In this setup, both the bead and tip where in the same imaging (focal) plane. If *B*_blunt_ was set to 0 Gs (i.e., *I* = 0 A), the bead would move towards the needle tip, evidence of a non-zero ***F***_MT_ secondary to permanent magnetization of the tip. By carefully adjusting *B*_blunt_, the field at the tip could be nullified (i.e., ***B***_tip_ could be forced to 0 Gs) and bead motion would stop because ***F***_MT_ = 0 nN. The value of *B*_blunt_ necessary to nullify ***B***_tip_ after degaussing the core is referred to as the permanent null flux density correction, defined here as 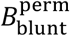. Values of 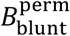 were observed to range from −2 to −4 Gs with a weak dependence on the needle core’s inclination angle (with respect to the horizontal). For an inclination angle of ~25° (standard MT configuration), 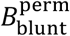 was −3.5 Gs.

Similarly, a corrective scheme was also engineered to mitigate the effects of the apparent remnant tip magnetization, similar to the coercive current corrections used in other MT devices^23^. To quantify these empirical corrections, the needle core was positioned in its standard configuration, and a 10-cycle OFF/ON 0.25-Hz waveform sequence was applied to the device using 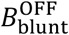 setpoints adjusted with the permanent null flux-density correction (i.e., 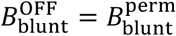). Here 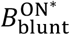 setpoints were used, spanning from 0 Gs to 185 Gs *relative* to the adjusted 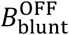, where 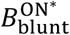 denotes the magnitude of 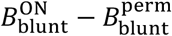. Following application of a given waveform sequence, the needle tip was positioned at δ = ~5 *μ*m from a superparamagnetic bead suspended in glycerol within the same focal plane. Despite the adjustment of 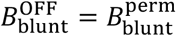, bead motion was still observed following magnetization of the core to all 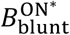 setpoints other than 0 Gs. However, bead motion could be stopped by further adjustment of 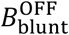, specifically, by setting 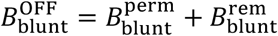, where 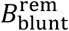 refers to the additional magnetic flux density correction required to nullify the apparent remnant field at ***B***_tip_. As shown in **Fig. 8**, 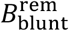 corrections were found to be a function of the 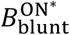 setpoint. Both sigmoidal and Langevin functions provide satisfactory fits of the data, and thereby a means for implementing an automated corrective algorithm in our overall *B*_blunt_-control scheme. Thus, by implementing the empirical null flux density corrections, 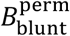 and 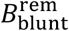, we were able to establish robust control of ***F***_MT_(*t*) in our MT device for applied square OFF/ON and multistep OFF/ON waveforms with 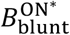 setpoints ranging from 0 to 185 Gs, frequencies ranging from 0.01 to 1 Hz, and 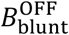 setpoints that effectively maintain ***F***_MT_ = 0 nN for all OFF intervals of the actuation waveform. Higher frequency actuation is possible but was not further explored in this work.

**FIG. 8.**
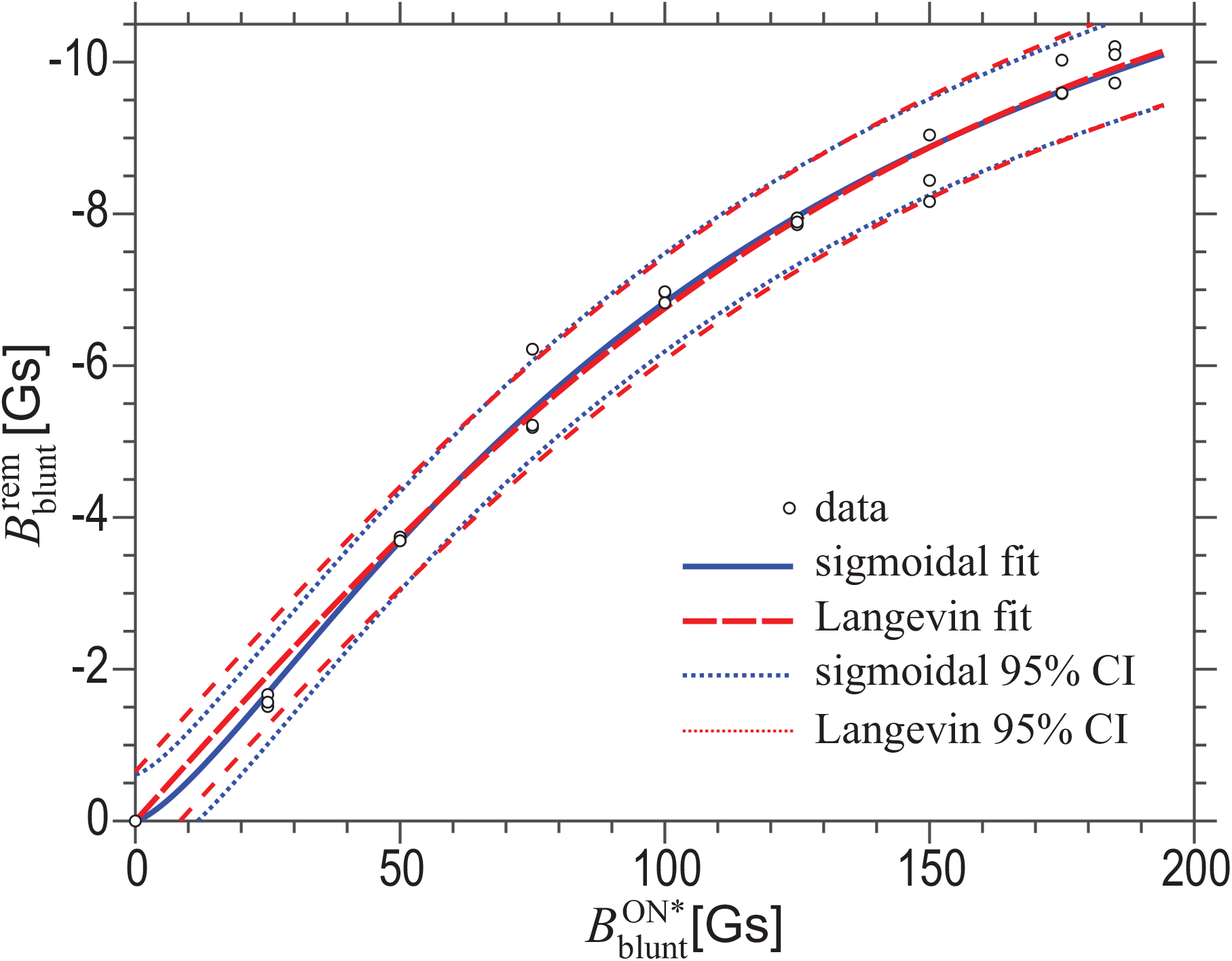
In addition to compensating for the permanent magnetization of the needle tip that exists despite degaussing of the core (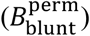), empiric corrections to 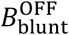 control points (defined here as 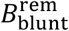) are also required to nullify ***B***_tip_ and hence achieve null magnetic actuation forces (***F***_MT_ = 0 nN) during the OFF portions of a prescribed actuation waveform sequence. Based on numerous empirical measurements, 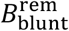 corrections were found to be a function of 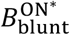 setpoints that can analytically be described by either a sigmoidal or Langevin function.

## V. CALIBRATION

Force calibration experiments were conducted in two different ranges: the *far* field of the needle tip (i.e., 50 *μ*m ≤ δ ≤ 200 *μ*m), and the *near* field of the tip (i.e., 2.5 *μ*m ≤ δ ≤ 50 *μ*m). Pure glycerol with a nominal dynamic viscosity of 1412 cP at 25°C (Sigma Aldrich) was used as the suspending medium for far field calibrations, and 10,000 cSt polydimethylsiloxane (PDMS) silicone oil (Sigma Aldrich) for near field calibrations, both subject to ambient environmental conditions of the lab (~22°C to 25°C and relative humidity of 10% to 50%). For both types of calibration, 4.5 *μ*m-diameter Dynabeads^®^ M-450 Tosylactivated superparamagnetic beads (Invitrogen) were suspended in their respective liquid media and dispensed onto 50 mm-diameter petri dishes with 35 mm-diameter No. 1.5 coverslip glass bottoms (MatTek). Bead concentration was dilute, such that individual beads were separated by at least 20-bead diameters to minimize interactions between beads during calibration^24^. Only beads at least 100 *μ*m above the surface of the coverslip were interrogated^24^. The MT device was placed in the standard configuration with an inclination angle of 25° to the horizon, and the tip was positioned a set distance away from a bead of interest, δ, via the three-axis micromanipulator while maintaining simultaneous focus of the bead and the needle tip under DIC imaging. A fast time-lapse DIC image acquisition (40 to 80 fps, depending on ROI settings) was then initiated using Nikon Elements, thereby triggering actuation of the MT device with a 0.25-Hz calibration OFF/ON waveform magnetic actuation sequence for a 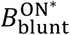 setpoint while mploying flux density corrections to achieve null magnetic actuation forces at the needle tip for all OFF setpoints (see <u>Sec. IV.E</u>). Calibrations were repeated for 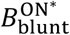 setpoints ranging from 0 Gs to 185 Gs.

Following each calibration experiment, the position of the centroid of the superparamagnetic bead and the position of the needle tip were tracked within each frame of the overall image sequence using a custom MATLAB code (Mathworks, Inc., Natick, MA) based on cross-correlation of sequential imaging frames^11^. Bead velocities, ***V***, and **δ** were evaluated for both OFF- and ON-portions of the MT actuation waveform sequence by temporally correlating DIC time-lapse images to magnetic flux density data using the recorded TTL exposure pulses output from the sCMOS camera. Data acquired during transient segments of the MT actuation waveform were omitted from the calibration analysis. To account for bead motion unrelated to magnetic actuation during the calibration experiment, *corrected* bead velocities during each ON interval of the waveform sequence, defined here as 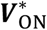, were calculated by subtracting the *average* bead velocity present in the immediately preceding OFF interval, 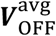, from the raw measurement of ***V***_ON_, or 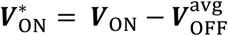 (see **Fig. 9**). During our calibration experiments, 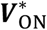 were observed to be oriented in a vectorially opposite direction to their corresponding **δ**. Moreover, the magnitudes of 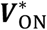 depended only on the magnitude of **δ** for beads positioned within a symmetric field in front of the needle, defined by a central axis consisting of the horizontal projection of the longitudinal axis of the needle core and a field bounded by lines extending outward from the needle tip at angles of approximately ±40° from this central axis^23,24^. Based on these experimental observations, knowing the dynamic viscosity of the suspending medium, η, and the diameter of the superparamagnetic bead, *d*, ***F***_MT_(δ) for the imposed 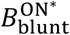 setpoint were calculated using the Stokes-Einstein equation^24,25^:

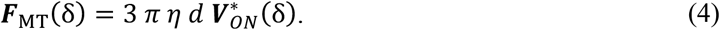

**FIG. 9.**
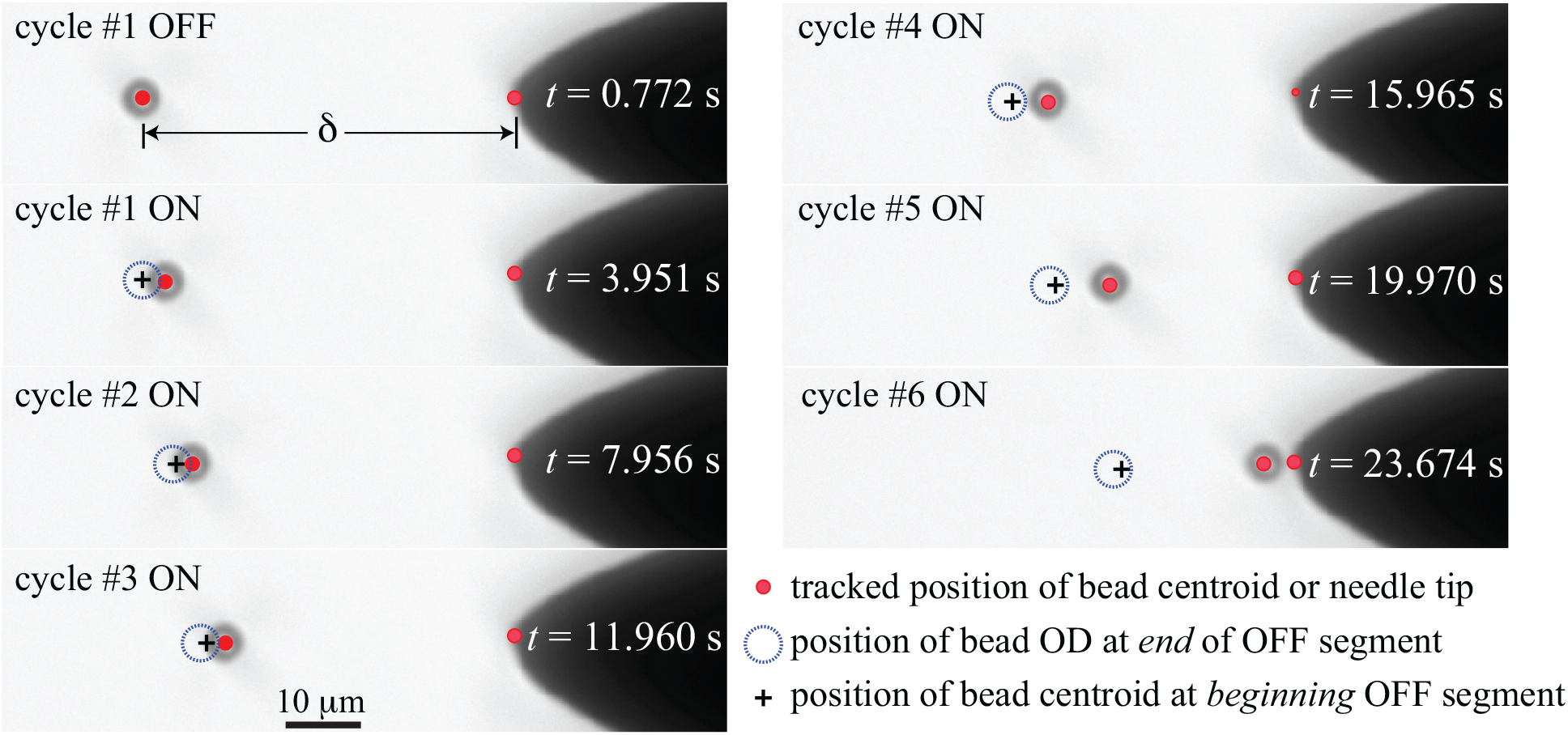
Force calibration of the MT device. A series of DIC images at select timepoints are shown demonstrating movement of a 4.5 μm-diameter superparamagnetic bead suspended in 10,000 cSt silicone oil and subject to actuation with the MT device using a 6-cycle OFF/ON 0.25 Hz *B*_blunt_-control waveform with 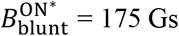. 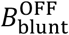 control setpoints were programmed to maintain null actuation forces during OFF portions of the calibration sequence. Time stamps correlate with the real-time movie of the calibration sequence that can be found here (**Vid 1**). During calibration, the bead drifts slightly *away* from the needle tip during the OFF segments of each actuation cycle, as visualized by the eccentricity between the bead’s tracked centroid at the beginning of each OFF segment (black cross) and the bead’s tracked outer diameter (OD) at the end of each OFF segment (blue dashed circle). Bead drift is unrelated to magnetic actuation.

Here, δ denotes the magnitude of the positional vector of the superparamagnetic bead, **δ**, calculated as the Euclidean separation distance between the centroid of the bead and the needle tip. Eq. (4) assumes low Reynolds numbers, i.e., equilibrium between ***F***_MT_(δ) and the drag force that a spherical object experiences under laminar flow conditions. Reynolds numbers for calibrations conducted in silicone oil and glycerol were ~10^−8^ and ~10^−7^, respectively. With this method of data reduction, a family of six discrete calibration δ-*F*_MT_ curves were generated, one for each 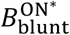 setpoint (where *F*_MT_ denotes the scalar magnitude of ***F***_MT_).

Near field calibrations conducted with silicone oil are shown in **Fig. 10**, while far field calibration experiments with glycerol are presented in **Fig. S3** in the **Supplementary Material**. As anticipated, for each δ-*F*_MT_ curve, *F*_MT_ exponentially increases with decreasing δ. For any given value of δ, *F*_MT_ increases with increasing 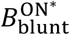 setpoint. Interestingly, data collected for 150 Gs, 175 Gs, and 185 Gs 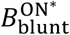 setpoints were experimentally indistinguishable, suggestive that ***B***_tip_ saturates before ***B***_blunt_ (see **Fig. 1(a)**). Arguably, this finding is consistent with the data shown in **Fig. 3**, where *B*_tip_ measurements demonstrated much more rapid saturation in magnetic flux density with increasing solenoid currents compared to *B*_blunt_. In other words, values of *B*_tip_ seem to exhibit little to no change as *B*_blunt_ ranges between 150 Gs and 185 Gs. Alternatively, we considered the possibility that the superparamagnetic beads were saturating in response to ***B***_tip_ for 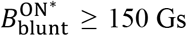 and thus leading to negligible increases *F*_MT_. However, M-450 beads are not known to exhibit magnetic saturation for applied fields of <1000 Gs, so this possibility seems unlikely^36^. The more plausible physical possibility is that the needle core overall exhibits a spatially non-uniform magnetization in response to an applied field, consistent with our previous observations of an apparent spatially non-uniform remnant magnetization.

**FIG. 10.**
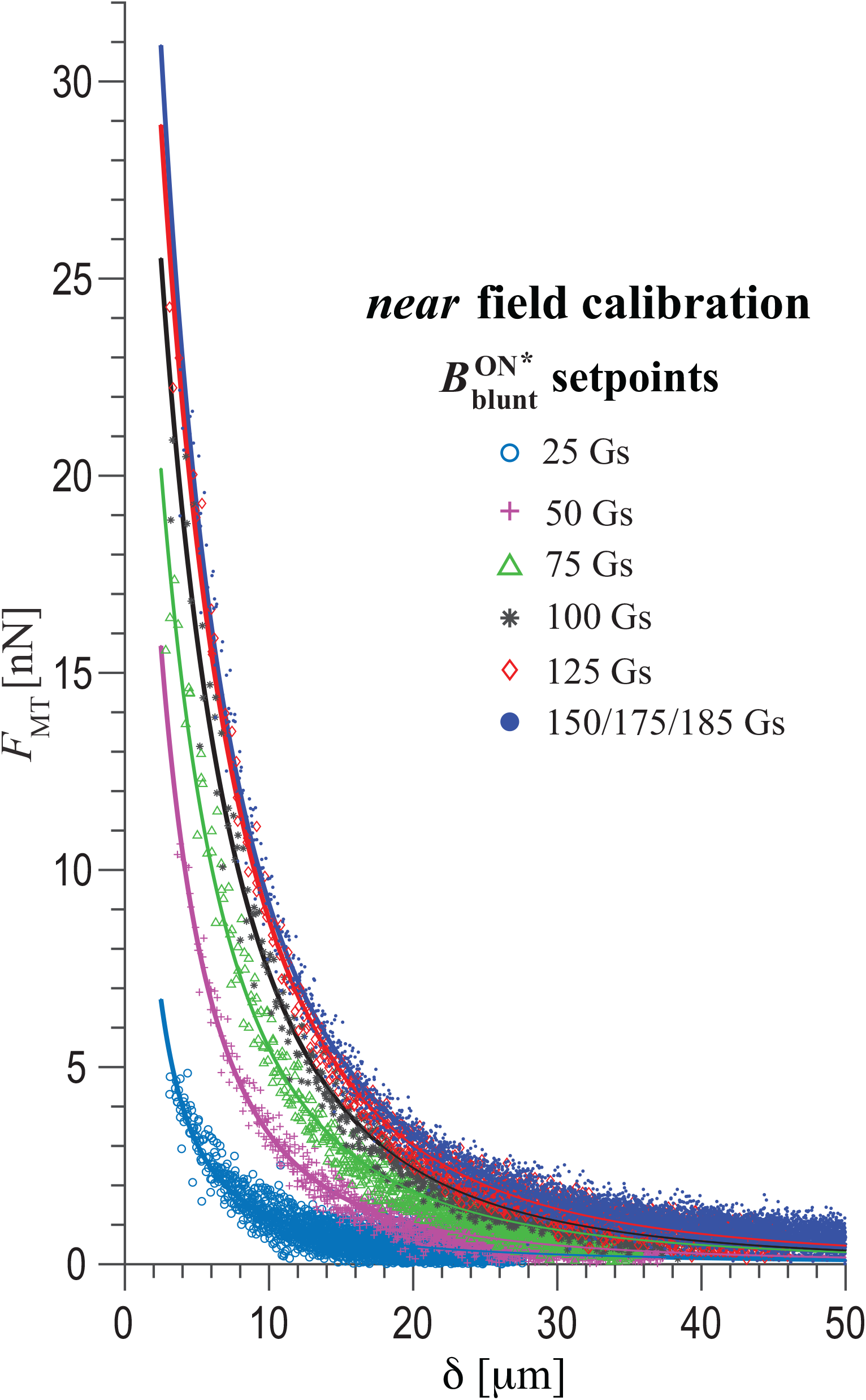
Near field calibration. A family of calibration curves that define *F*_MT_(δ) over the domain 2.5 *μ*m ≤ δ ≤ 50 *μ*m are shown where solid lines represent the best-fit three-term power law given by **Eq. (5)** for each respective data set.

Calibrations conducted with silicone oil exhibited minor differences in *F*_MT_ predictions when subject to small variations in room temperature and relative humidity that were present within the laboratory environment. In contrast, large variations in *F*_MT_ for the same 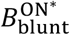 flux density setpoint appeared for calibrations conducted in glycerol when deviations in ambient temperature and relative humidity were not accounted for in the assumed value of η (see **Fig. 11**). Due to its hygroscopic nature, a thin film (<1000 μm) of pure glycerol exposed to laboratory conditions rapidly equilibrates with water vapor present in the ambient air forming a water-glycerol mixture. Consequently, the dynamic viscosity of this equilibrium glycerol-water mixture is a function of both relative humidity and temperature^37–39^. For example, at 20°C, viscosities of glycerol-water mixtures vary more than 12-fold from 765 cP to 60.1 cP as the relative humidity varies from 10% to 50%^38,40^. Similarly, though of smaller magnitude, A temperature increase from 20° to 30°C reduces the dynamic viscosity by ~50%^39^. **Fig. 11** compares a 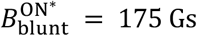 calibration conducted in silicone oil at 25°C to a calibration done in glycerol at 25°C. Here, the calibrations assume a dynamic viscosity for either pure glycerol based on a relative humidity of 0% or a water-glycerol mixture at 25°C that accounts for a relative humidity of 19% present in the ambient air. When humidity is properly accounted for, the silicone and glycerol calibrations demonstrate excellent quantitative agreement. However, glycerol calibrations that do not account for humidity effects generate erroneous *F*_MT_ values, with several-fold deviations in predicted magnetic actuation forces in the near field when compared to calibrations conducted in silicone. With this insight, it seems plausible that some of the discrepancy in force capabilities of previously described MT devices might be secondary to glycerol-based calibrations done in the absence of temperature and humidity assessments.

**FIG. 11.**
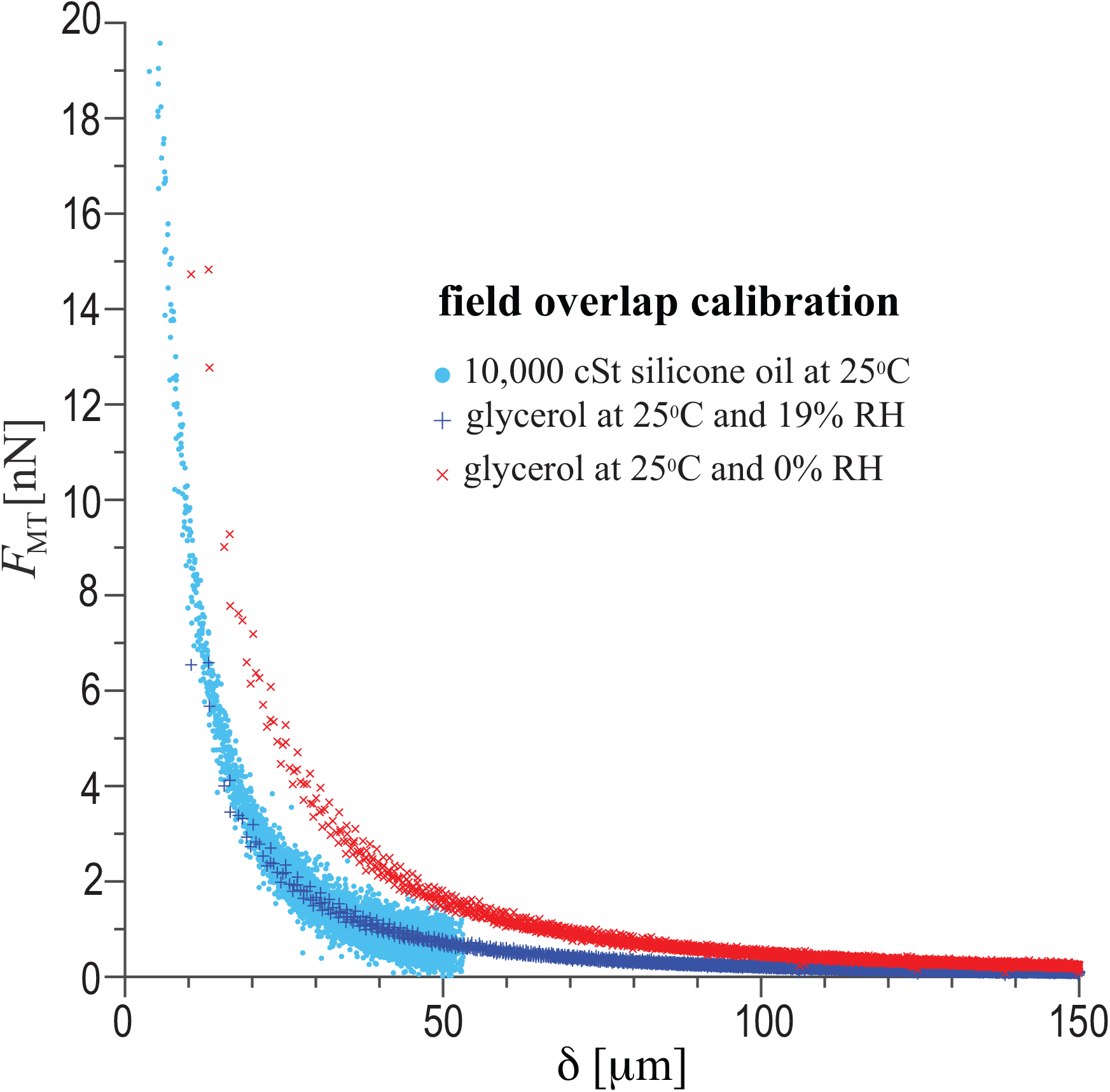
Quantitative overlap between near field and far field calibration data requires accounting for the relative humidity-dependent effects on the dynamic viscosity of glycerol.

Fits of near field *F*_MT_(δ) calibration measurements were done for each of the six 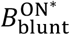 flux density setpoints using the following three-term power law^33^:

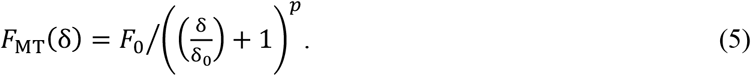

Fit parameters for each calibration curve are shown in **Table I**. Fit of the 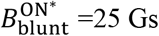 was poor. Values of *F*_0_ increase with increasing magnitudes of 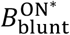 setpoints. With the exception of the 25 Gs 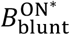 setpoint, *p* ranged from ~2.3 to 2.9, consistent with the modeling predictions of Bijamov et al.^33^ Moreover, we note that the best-fit values of sδ_0_ are ~10 μm, a value that is roughly equivalent to the radius of curvature of our needle tip, a finding also observed by Bijamov et al.^33^ We attempted to assign a universal fit for all of the data as done by Kollmannsberger and Fabry^23^, but this led to unacceptable uncertainties in *F*_MT_(δ) predictions. Consequently, for our MT device, *F*_MT_(δ) are evaluated by means of three-parameter power-law fits specific to each of the six calibrated 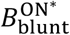 flux density setpoints. Using a standard approach to uncertainty analysis^41^, defining Δ*F*_MT_(δ) as the uncertainty in *F*_MT_(δ) for a 95% confidence level, and assuming ±1 *μ*m for the uncertainty in δ at a 95% confidence level for experiments in which the needle tip and the superparamagnetic bead are both present within the same horizontal imaging plane, the relative uncertainty, ω*F*_MT_, defined here as Δ*F*_MT_(δ) /*F*_MT_(δ), for various 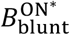 setpoints and for δ spanning from 2.5 *μ*m to 50 *μ*m are plotted in **Fig. 12**. Excluding the 25 Gs, 50 Gs, and 75 Gs 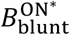 setpoints, relative uncertainties in *F*_MT_(δ) are <30% for 2.5 μm < δ < 40 μm, when using setpoints ≥100 Gs. Note that small variations of superparamagnetic bead diameter are accounted for within the uncertainties of the power law fit parameters because each calibration curve was based on testing of at least 5 independent beads. Overall, our *F*_MT_(δ) uncertainties are quantitatively similar to those reported by Kollmansburger et al.^23^

**TABLE 1.** Power law fits.

| $B_{\text{blunt}}^{\text{ON}^*}$ [Gs] | $F_0$[nN] | $p$ | $\delta_0$[$\mu\text{m}$] | $F_0$ [nN]<br>95% CI | $p$ 95% CI | $\delta_0$ [$\mu\text{m}$]<br>95% CI | $R^2$ |
| --- | --- | --- | --- | --- | --- | --- | --- |
| 25 | 35.46 | 1.568 | 1.323 | $\pm 13.72$ | $\pm 0.090$ | $\pm 0.558$ | 0.7744 |
| 50 | 38.30 | 2.312 | 5.304 | $\pm 3.766$ | $\pm 0.114$ | $\pm 0.309$ | 0.9355 |
| 75 | 38.52 | 2.508 | 8.503 | $\pm 2.515$ | $\pm 0.150$ | $\pm 1.164$ | 0.9286 |
| 100 | 44.92 | 2.923 | 11.709 | $\pm 1.841$ | $\pm 0.143$ | $\pm 1.147$ | 0.9764 |
| 125 | 50.07 | 2.772 | 11.401 | $\pm 1.570$ | $\pm 0.098$ | $\pm 0.836$ | 0.9826 |
| 150/175/185 | 58.65 | 2.331 | 8.206 | $\pm 1.322$ | $\pm 0.036$ | $\pm 0.334$ | 0.9759 |

CI = confidence interval

**FIG. 12.**
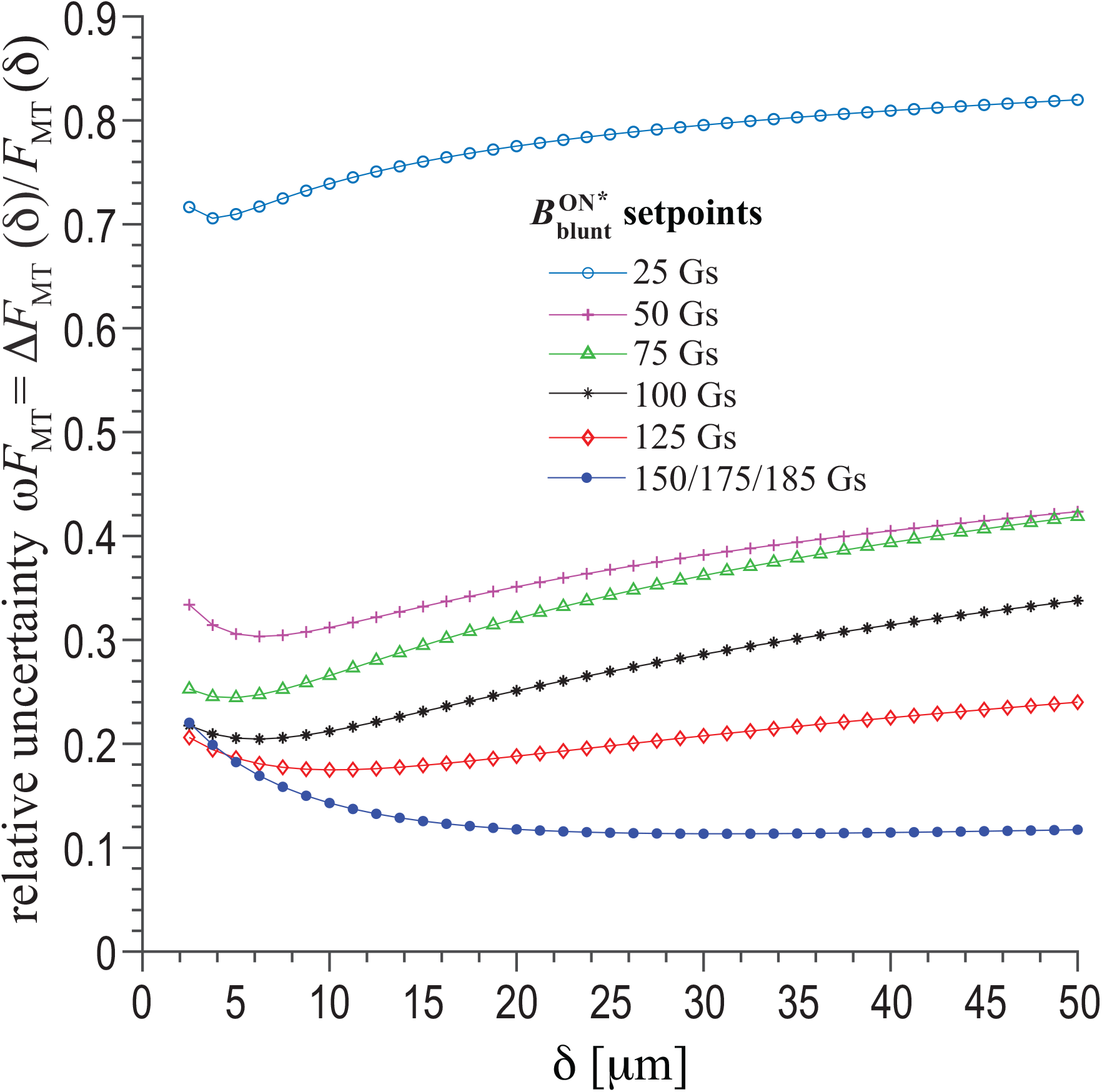
Relative uncertainties of magnetic actuation force within the near field, defined here as ω*F*_MT_(δ), as a function of 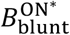, with *F*_MT_(δ) calculated using **Eq. (5)** following a standard uncertainty analysis at a 95% confidence level.

## VI. CONCLUSIONS

As motivation for this work, we propose that a magnetic tweezers (MT) device, integrated with substrate deformation tracking microscopy (DTM) and traction force microscopy (TFM), can be used to investigate the biophysical mechanisms of human blistering skin disease. While integration and application of MT with DTM/TFM is presented in a companion paper^20^, here we have presented key findings on the design, assembly, operation, and calibration of an MT device that is amenable to replication in any biology laboratory. Unique to our device, we have employed feedback control of the magnetic flux density emanating from the blunt end of the soft ferromagnetic needle core as an alternative to conventional devices that are based on feedback control of solenoid current. Using a cascade control scheme with PID gain scheduling, loading and unloading response times of our device (<100 ms) are necessarily slower than conventional current-based feedback-controlled MT devices. However, by intentionally avoiding overshoot and by intrinsically compensating for thermal heating effects, magnetic flux density-based feedback control of an MT device achieves more reproducible magnetic fields at the needle tip and are hence less cycle-to-cycle variation in magnetic actuation forces. Direct measurements of magnetic fields emanating from the needle tip suggest the existence of a small but finite permanent magnetization of the tip that remains even after degaussing of the needle core. Additionally, our observations suggest that the needle core experiences a spatially non-uniform magnetization in response to the applied field of the solenoid coil, where fields at the tip demonstrate more rapid saturation and increased remnant magnetization compared to fields emanating from the blunt end of the core. Further investigation will be required to determine the physical mechanisms that give rise to non-uniform magnetization of the core. Nevertheless, empirically derived control schemes can be developed to achieve null magnetic actuation forces and robust operation of an MT device based on magnetic flux density control. Rigorous calibration of our MT device has validated its ability to produce magnetic actuation forces up to 25 nN on 4.5 *μ*m-diameter superparamagnetic beads with a maximum relative error of ±30% using magnetic flux density setpoints ≥100 Gs and for beads positioned between 2.5 *μ*m and 40 *μ*m from the needle tip.

## Supporting information

Vid 1 calibration

## VII. SUPPLEMENTARY MATERIAL

A comprehensive list of abbreviations and symbols used in this work, photographs of the magnetic tweezers instrumentation and setup, and far field calibration data can be found in the **Supplementary Material**.

## VIII. ACKNOWLEDGEMENTS

The authors acknowledge Dan Witte and George DeBeck (Nikon Instruments, Inc.) for their much-needed technical assistance with the setup and operation of our microscope. E.A.S. thanks the National Science Foundation (CAREER 1452728). J.C.S. thanks the Dermatology Foundation for their support of this work through a career development award.

## IX. AUTHOR’S CONTRIBUTIONS

All authors contributed equally to this work.

## X. DATA AVAILABILITY

The data that supports the findings of this study are available within the article and its supplementary material. Electronic copies of the LabVIEW™ virtual instruments (VIs) used to control the MT device are available upon reasonable request.

