## Supplementary Material for "Magnetic Tweezers with Magnetic Flux Density Feedback Control"

### LIST OF ABBREVIATIONS AND SYMBOLS

| | |
|---|---|
| $\boldsymbol{B}_{\text{blunt}}$ | magnetic field at the blunt end of the needle |
| $B_{\text{blunt}}$ | magnetic flux density emanating in the axial direction from the blunt end of the needle core |
| $B_{\text{blunt}}^{\text{ON}}$ | $B_{\text{blunt}}$ setpoint during the ON portion of a magnetic actuation waveform |
| $B_{\text{blunt}}^{\text{ON}*}$ | $B_{\text{blunt}}^{\text{ON}}$ setpoint relative to $B_{\text{blunt}}^{\text{perm}}$ |
| $B_{\text{blunt}}^{\text{OFF}}$ | $B_{\text{blunt}}$ setpoint during the OFF portion of a magnetic actuation waveform (null magnetic force setpoint) |
| $B_{\text{blunt}}^{\text{perm}}$ | $B_{\text{blunt}}$ setpoint necessary to nullify $\boldsymbol{B}_{\text{tip}}$ following degaussing of the needle core |
| $B_{\text{blunt}}^{\text{rem}}$ | additional magnetic flux density setpoint necessary to nullify $\boldsymbol{B}_{\textbf{tip}}$ following magnetization of the needle core |
| $\Delta B_{\text{blunt}}^{\text{ON}}$ | difference in the time-averaged ON setpoints observed between cycle #10 and cycle #1, as measured by $B_{\text{blunt}}$ |
| $\Delta B_{\text{blunt}}^{\text{OFF}}$ | difference in the time-averaged OFF setpoints observed between cycle #10 and cycle #1, as measured by $B_{\text{blunt}}$ |
| $\boldsymbol{B}_c$ | magnetic field within the needle core |
| $B_c^{\text{sat}}(T)$ | temperature-dependent saturation magnetization of the needle core |
| $B_c^{\text{rem}}(T)$ | temperature-dependent remnant magnetization within the needle core |
| $\boldsymbol{B}_{\text{tip}}$ | magnetic field at the needle tip |
| $B_{\text{tip}}$ | magnetic flux density measured within the field of the needle tip using a vertically oriented Hall effect sensor with the center of its sensing loop positioned lateral to the needle tip within the same horizontal plane |
| $\Delta B_{\text{tip}}^{\text{ON}}$ | difference in the time-averaged ON setpoints observed between cycle #10 and cycle #1, as measured by $B_{\text{tip}}$ |
| $\Delta B_{\text{tip}}^{\text{OFF}}$ | difference in the time-averaged OFF setpoints observed between cycle #10 and cycle #1, as measured by $B_{\text{tip}}$ |

| | |
|---|---|
| $d$ | superparamagnetic bead diameter |
| $\boldsymbol{\delta}$ | spatial positioning vector of superparamagnetic bead with respect to the needle tip |
| $\delta$ | scalar magnitude of the spatial positioning vector of superparamagnetic bead with respect to the needle tip, $\boldsymbol{\delta}$ |
| $\delta_0$ | distance term in power law describing $F_{\mathrm{MT}}$ (see **Eq. (5)** of the main article) |
| $\boldsymbol{F}_{\mathrm{MT}}$ | magnetic actuation force |
| $F_{\mathrm{MT}}$ | scalar magnitude of magnetic actuation force |
| $\Delta F_{\mathrm{MT}}(\delta)$ | uncertainty in $F_{\mathrm{MT}}(\delta)$ as a function of $\delta$ (95% confidence level) |
| $\omega F_{\mathrm{MT}}$ | relative uncertainty in $F_{\mathrm{MT}}(\delta)$ or $\Delta F_{\mathrm{MT}}(\delta)\ /F_{\mathrm{MT}}(\delta)$ |
| $F_0$ | force term in power law describing $F_{\mathrm{MT}}$ (see **Eq. (5)** of the main article) |
| $\eta$ | dynamic viscosity of liquid medium |
| $H_c$ | coercivity of the needle core |
| $I$ | solenoid current |
| $I(t)_{\mathrm{dmag}}$ | degaussing solenoid current |
| $I_0$ | initial current setting for degaussing solenoid current |
| $\mu_b$ | relative permeability of superparamagnetic bead |
| $\mu_c(T)$ | temperature-dependent relative permeability of the needle core |
| $\mu_0$ | permeability of free space |
| $p$ | exponent in power law describing $F_{\mathrm{MT}}$ (see **Eq. (5)** of the main article) |
| $\boldsymbol{V}_{\mathrm{ON}}$ | raw measurement of superparamagnetic bead velocity during the ON interval of a magnetic actuation waveform |
| $\boldsymbol{V}_{\mathrm{OFF}}^{\mathrm{avg}}$ | average superparamagnetic bead velocity measured during the preceding OFF interval of a magnetic actuation waveform |

| | |
|---|---|
| $\boldsymbol{V}_{\mathrm{ON}}^{*}$ | superparamagnetic bead velocity corrected for motion unrelated to magnetic actuation |
| $T$ | temperature |
| $t$ | time |

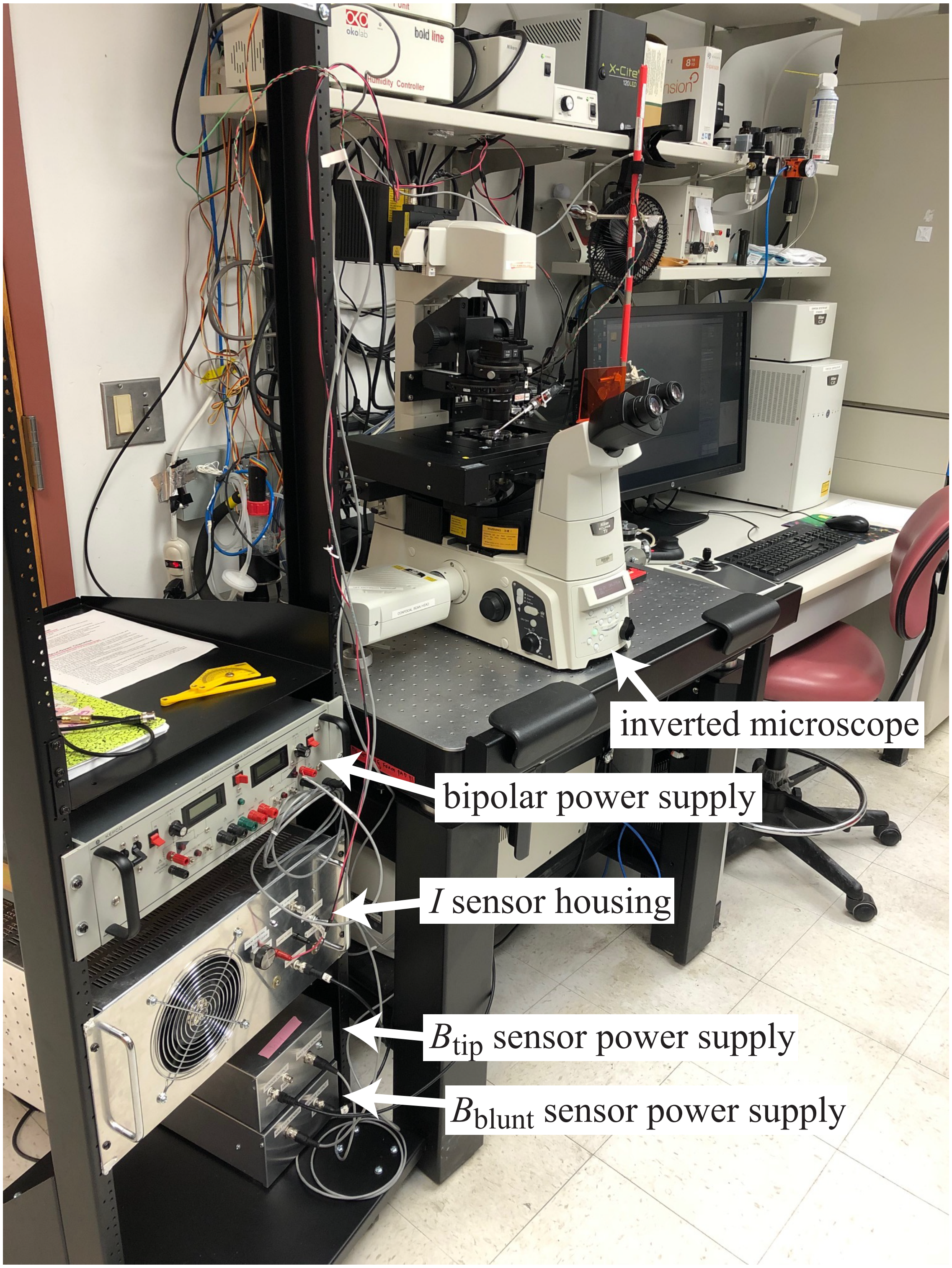


**FIG. S1.** Major MT device components.

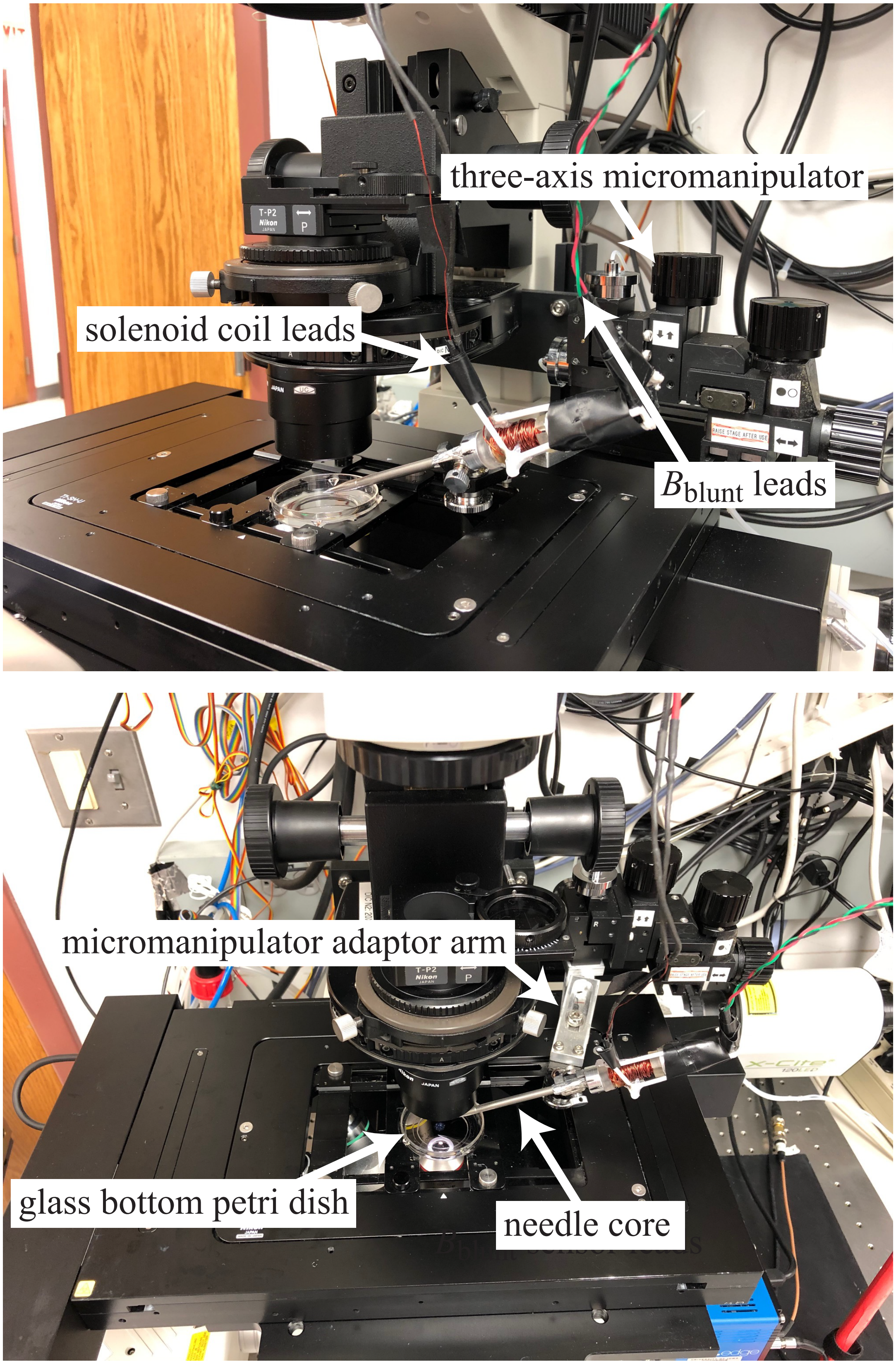


**FIG. S2.** MT device components on the microscope stage.

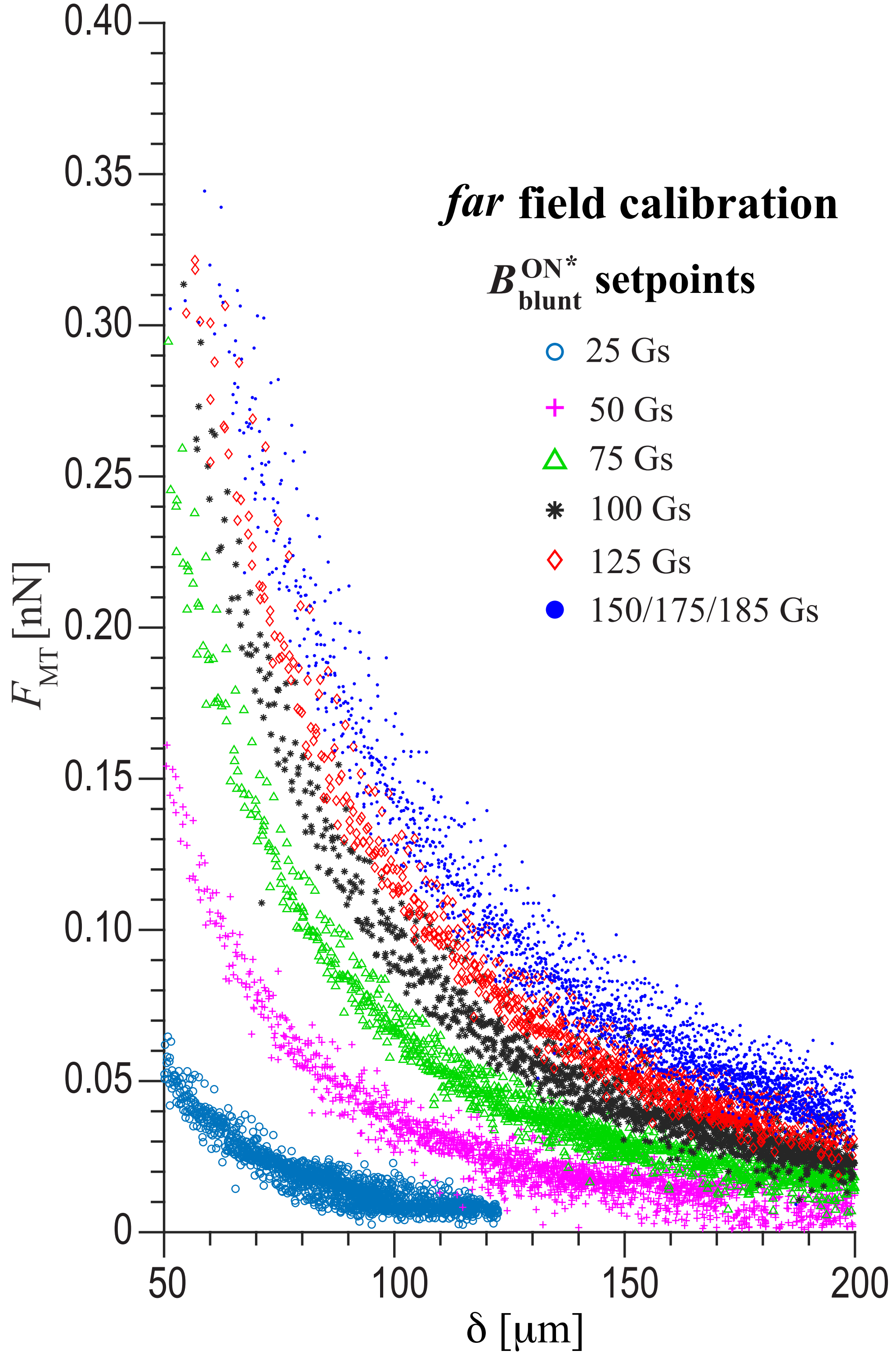


**FIG. S3.** Far field calibration data collected by while using the MT device to magnetically actuate 4.5 μm-diameter superparamagnetic beads suspended in glycerol. Power law fits were not done for these data sets.
